# The previously uncharacterized RnpM (YlxR) protein modulates the activity of ribonuclease P in *Bacillus subtilis*

**DOI:** 10.1101/2023.07.28.550789

**Authors:** Dennis Wicke, Piotr Neumann, Markus Gößringer, Aleksandar Chernev, Anja Poehlein, Rolf Daniel, Henning Urlaub, Roland K. Hartmann, Ralf Ficner, Jörg Stülke

## Abstract

Even though *Bacillus subtilis* is one of the most studied organisms, no function has been identified for about 20% of its proteins. Among these unknown proteins are several RNA- and ribosome-binding proteins suggesting that they exert functions in cellular information processing. In this work, we have investigated the RNA-binding protein YlxR. This protein is widely conserved in bacteria and strongly constitutively expressed in *B. subtilis* suggesting an important function. We have identified the RNA subunit of the essential RNase P as the binding partner of YlxR. The main activity of RNase P is the processing of 5’ ends of pre-tRNAs. *In vitro* processing assays demonstrated that the presence of YlxR results in reduced RNase P activity. Chemical cross-linking studies followed by *in silico* docking analysis and experiments with site-directed mutant proteins suggest that YlxR binds to the region of the RNase P RNA that is important for binding and cleavage of the pre-tRNA substrate. We conclude that the YlxR protein is a novel interaction partner of the RNA subunit of RNase P that serves to finetune RNase P activity to ensure appropriate amounts of mature tRNAs for translation. We rename the YlxR protein RnpM for RNase P modulator.

## INTRODUCTION

The Gram-positive bacterium *Bacillus subtilis* is one of the most intensively studied and best understood organisms. The interest in this bacterium was triggered by its ability to sporulate, *i.e*. to undergo a simple developmental program that results in a dormant state allowing long-term survival under adverse conditions. Moreover, *B. subtilis* is an important workhorse in biotechnology, *i.e.* for the production of vitamins and enzymes, and it is a close relative of many important pathogens, such as *Listeria monocytogenes* and *Staphylococcus aureus* (1). The genome of *B. subtilis* was one of the first to be fully sequenced; however, of its about 4,200 protein-coding genes, nearly 20% or around 800 proteins still await their functional assignment (2).

The fact that no function is known for about 800 proteins of *B. subtilis* indicates that there are large unexplored areas that require intensive research. However, most of the proteins of unknown functions are probably required only under very specific conditions. These unknown proteins that account for about 20% of all proteins need only 2.7% of the translation capacity for their synthesis, indicating that they are strongly underrepresented in the *B. subtilis* proteome (3). Indeed, most of the unknown proteins are very poorly expressed under most conditions; however, a subset of unknown proteins is highly expressed under more than 100 different conditions (4). While the investigation of poorly expressed proteins may appear only of limited relevance for the understanding of bacterial physiology under standard laboratory conditions, it is an important challenge to analyze the functions of the set of unknown proteins that are highly expressed and likely exert key functions in the physiology and/or cell biology of *B. subtilis* (5).

We are interested in gaining a comprehensive understanding of the cellular activities of the model bacterium *B. subtilis*. As stated above, the investigation of highly expressed proteins of unknown function is of particular importance in this context. We have compiled a list of 40 highly expressed genes of unknown functions that we consider as being of interest as primary research targets (5). Many of these proteins are thought to bind RNA and/ or the ribosome and may thus have vital functions in molecular information processing in the cell.

Our group has a long-standing interest in studying protein-RNA interactions (6, 7, 8, 9, 10), and we have been particularly intrigued by the identification of KhpA and YlxR, two highly expressed unknown proteins, that were observed to interact with RNA in other bacteria (11, 12). Both proteins are widespread among bacteria, with the notable exception of Beta- and Gammaproteobacteria (and also Alphaproteobacteria in the case of KhpA) (5). The KhpA protein consists of a KH domain that is found in a variety of RNA-binding enzymes as well as other proteins (12, 13). KhpA has been implicated in cell division in *Streptococcus pneumoniae* (14); however, the precise mechanism by which the protein would affect cell division is unknown. Interestingly, the Hfq protein that facilitates RNA-RNA interactions in *E. coli* (15), is either missing in many Bacilli or, if present, did not show a role in mediating RNA-RNA interactions (16, 17, 18) suggesting that this protein has a different function in those Bacilli that encode a Hfq paralog. It has thus been proposed that KhpA, rather than Hfq, might act as an RNA matchmaker in Gram-positive bacteria (12).

The YlxR protein shares structural features with RNA-binding proteins, such as a large positively charged surface patch on one side of the protein comprising a large cleft (19). Based on the structure, it was already suggested that YlxR might be an RNA-binding protein, a possibility that recently gained support from a global analysis of RNA-binding proteins in *Clostrioides difficile* (11). The *ylxR* gene is part of the *rimP-nusA-ylxR-rplGA-infB* operon that is strongly conserved in Firmicutes, Actinobacteria, and Cyanobacteria. All genes of this operon play important roles in transcription (*nusA*), ribosome maturation (*rimP*), or translation (*rplGA*, *infB*) suggesting that YlxR could also have a function related to these processes. Loss of the YlxR protein results in altered expression of a large set of genes with a wide array of diverse functions (20, 21).

One essential function in all organisms is the maturation of tRNAs. These molecules are synthesized as precursors, and are processed by RNases P and Z, respectively, at their 5’ and 3’ ends (22). Many tRNAs are initially synthesized without the essential CCA trinucleotide at the 3’ end of the processed molecule, in these cases the CCA is added by a dedicated tRNA nucleotidyltransferase (23). All three enzymes are essential in *B. subtilis* and many other organisms. Bacterial RNase P is typically composed of a catalytically active RNA subunit (the P RNA or RnpB RNA, encoded by the *rnpB* gene) and of a single protein cofactor (termed P protein or RnpA, encoded by *rnpA*), thus representing a minimal ribonucleoprotein complex. In this complex, the RNA molecule exhibits the catalytic activity, and the protein binds close to the active site of the P RNA to facilitate substrate recognition and catalysis under physiological conditions (24). So far, no mechanism for the regulation of RNase P activity has been identified in *B. subtilis* or any other bacteria.

The identification of functions of highly expressed proteins of unknown function is important to get a full understanding of the physiology, molecular and cell biology of any organism. In this study, we have addressed the functions of the potentially RNA-binding proteins KhpA and YlxR in *B. subtilis*. While no interacting RNA could be identified for KhpA, we found YlxR to bind to the P RNA, the essential RNA component of RNase P. We demonstrate that binding of YlxR to the P RNA results in a modest inhibition of RNase P activity. Chemical crosslinking, structural modeling of the ribonucleoprotein complex, and site-directed mutagenesis suggest that YlxR binds close to the active site of RNase P thus explaining the observed inhibition. Based on our results, we rename YlxR RnpM (<u>RN</u>ase <u>P m</u>odulator). This designation will be used hereafter.

## MATERIAL AND METHODS

### Bacterial strains, plasmids and growth conditions

All *B. subtilis* strains used in this study are derived from the laboratory strain 168 (*trpC2*). *B. subtilis* and *E. coli* cells were grown in Lysogeny Broth (LB medium) (25). LB plates were prepared by addition of 17 g Bacto agar/l (Difco) to LB (25).

### DNA manipulation

*B. subtilis* was transformed with plasmids, genomic DNA or PCR products according to the two-step protocol (25, 26). Transformants were selected on LB plates containing erythromycin (2 µg/ml) plus lincomycin (25 µg/ml), chloramphenicol (5 µg/ml), kanamycin (10 µg/ml), or spectinomycin (250 µg/ml). Competent cells of *E. coli* were prepared and transformed following the standard procedure (25) and selected on LB plates containing kanamycin (50 µg/ml). S7 Fusion DNA polymerase (Biozym Scientific GmbH, Germany) was used as recommended by the manufacturer. DNA fragments were purified using the QIAquick PCR Purification Kit (Qiagen, Germany). DNA sequences were determined by the dideoxy chain termination method (25). Chromosomal DNA from *B. subtilis* was isolated using the peqGOLD Bacterial DNA Kit (Peqlab, Erlangen, Germany).

### Construction of deletion mutants

Deletion of the *csrA, khpA, ptsH,* and *rnpM* (*ylxR*) genes was achieved by transformation with PCR products constructed using oligonucleotides to amplify DNA fragments flanking the target genes and intervening antibiotic resistance cassettes as described previously (27, 28, 29). The identity of the modified genomic regions was verified by DNA sequencing. The resulting strains were GP469, GP3752, GP3751, and GP3753, respectively.

### Plasmids

All plasmids used in this work are listed in Table 1. The plasmids pBQ200 (30), pGP380 and pGP382 (31) were used to construct expression vectors. For this purpose, the genes of interest were amplified by PCR using appropriate primer pairs that add restriction sites to the fragments. For *csrA*, the PCR product was cloned between the BamHI and PstI sites of pGP381. The resulting plasmid was pGP381. The *ptsH* gene was amplified with primers that also added the coding sequence for the Strep-tag and cloned between the BamHI and HindIII sites of pBQ200. This resulted in plasmid pG961. The *khpA* and *rnpM* amplicons were cloned between the XbaI and PstI sites of pGP382, yielding pGP3616 and pGP3617, respectively.

**Table 1.**
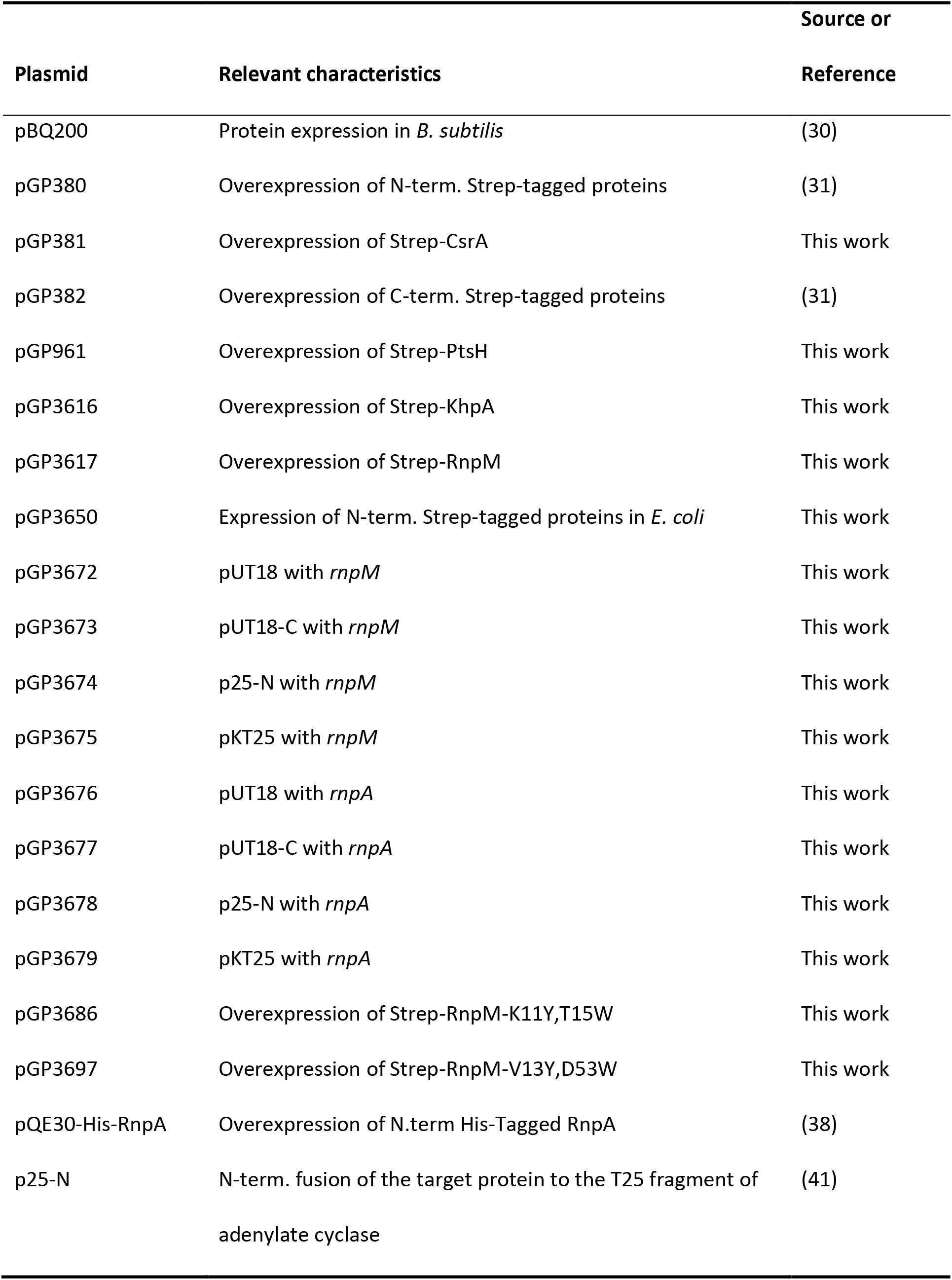
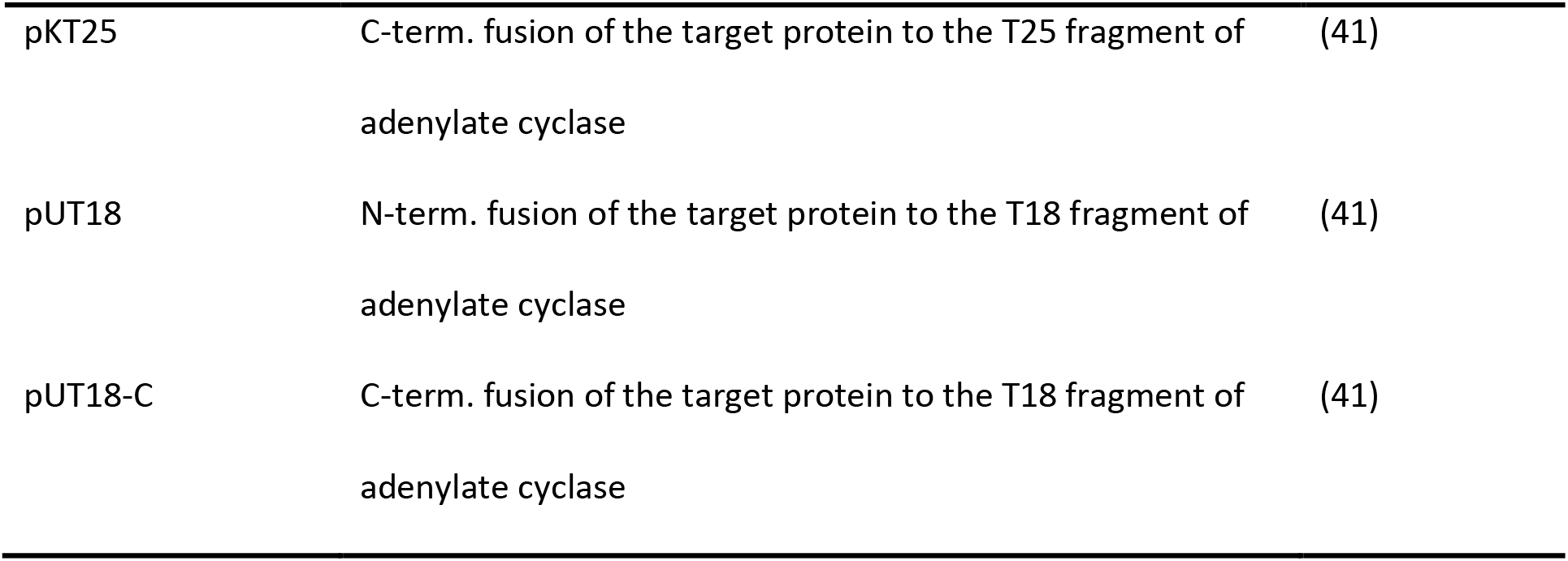
Plasmids used in this study.

| Plasmid | Relevant characteristics | Source or<br>Reference |
| --- | --- | --- |
| pBQ200 | Protein expression in *B. subtilis* | (30) |
| pGP380 | Overexpression of N-term. Strep-tagged proteins | (31) |
| pGP381 | Overexpression of Strep-CsrA | This work |
| pGP382 | Overexpression of C-term. Strep-tagged proteins | (31) |
| pGP961 | Overexpression of Strep-PtsH | This work |
| pGP3616 | Overexpression of Strep-KhpA | This work |
| pGP3617 | Overexpression of Strep-RnpM | This work |
| pGP3650 | Expression of N-term. Strep-tagged proteins in *E. coli* | This work |
| pGP3672 | pUT18 with *rnpM* | This work |
| pGP3673 | pUT18-C with *rnpM* | This work |
| pGP3674 | p25-N with *rnpM* | This work |
| pGP3675 | pKT25 with *rnpM* | This work |
| pGP3676 | pUT18 with *rnpA* | This work |
| pGP3677 | pUT18-C with *rnpA* | This work |
| pGP3678 | p25-N with *rnpA* | This work |
| pGP3679 | pKT25 with *rnpA* | This work |
| pGP3686 | Overexpression of Strep-RnpM-K11Y,T15W | This work |
| pGP3697 | Overexpression of Strep-RnpM-V13Y,D53W | This work |
| pQE30-His-RnpA | Overexpression of N.term His-Tagged RnpA | (38) |
| p25-N | N-term. fusion of the target protein to the T25 fragment of<br>adenylate cyclase | (41) |
| pKT25 | C-term. fusion of the target protein to the T25 fragment of<br>adenylate cyclase | (41) |
| pUT18 | N-term. fusion of the target protein to the T18 fragment of<br>adenylate cyclase | (41) |
| pUT18-C | C-term. fusion of the target protein to the T18 fragment of<br>adenylate cyclase | (41) |

To express RnpM mutant proteins for the final identification of the P RNA interaction area, the plasmids pGP3686 and pGP3697 were constructed to fuse a Strep-tag to the C-terminus of RnpM^K11Y,T15W^ and RnpM^V13Y,D53W^, respectively. RnpM variants were obtained by combined-chain reaction (CCR, 32) using pGP3617 as a template and the primer pair DW17/DW18 in combination with the mutagenesis primers DW180 for RnpM^K11Y,T15W^ and DW183 and DW184 for RnpM^V13Y,D53W^.

For the recombinant expression of Strep-tagged RnpM in *E. coli*, the plasmids pGP3650 and pGP3651 were constructed to fuse a Strep-tag to the N- or C-terminus of RnpM using the plasmids pGP172 (33) and pGP574 (34), respectively. The *rnpM* coding sequence was amplified using chromosomal DNA of *B. subtilis* 168 as the template and the primer pairs DW131/DW132 and DW131/DW133 for pGP3650 and pGP3651, respectively. The resulting fragments were cloned between the *Sac*I and *Kpn*I sites of pGP172 for the construction of pGP3650 and between *Sac*I and *Bam*HI sites of pGP574 for the construction of pGP3651.

To express and purify tag-free RnpM protein, *rnpM* was amplified using the primer pair DW40/DW41 and cloned between the *Bsa*I and *Xho*I sites of the expression vector pETSUMOadapt (Invitrogen, Germany). The resulting plasmid was pGP3618.

### Protein purification

His-SUMO-tagged RnpM was purified from *E. coli* Rosetta (DE3) carrying plasmid pGP3618. Protein expression was induced by the addition of isopropyl 1-thio-ß-galactopyranoside (IPTG, final concentration, 1 mM). His-SUMO-tagged RnpM was purified using 1 × ZAP Puffer (50 mM Tris-HCl, 200 mM NaCl, pH 7.5). Cells were lysed by four passages at 18,000 p.s.i. through an HTU DIGI-F press (G. Heinemann, Germany). After lysis, the crude extract was centrifuged at 35,000 × g for 45 min and then passed over a Ni^2+^nitrilotriacetic acid column (IBA, Germany). To remove attached RNA, 8 column volumes of 4 M LiCl were applied to the column. The protein was eluted with an imidazole gradient. After elution, the fractions were tested for the desired protein using SDS-PAGE. To remove the SUMO tag from the protein, the relevant fractions were combined, and the SUMO tag was removed by adding the SUMO protease (ratio 100:1) during overnight dialysis against 1 × ZAP buffer. The tag-free RnpM protein was collected and concentrated using a Vivaspin turbo 15 (Sartorius, Germany).

For expression and purification of Strep-tagged proteins, *E. coli* BL21(DE3) carrying the relevant plasmids was used. Cleared lysates of cells expressing the desired proteins were prepared as described (35) and subsequently passed over columns containing 1 ml Strep-Tactin matrix (IBA, Germany). The columns were washed with buffer W (100 mM Tris/HCl pH 8.0, 150 mM NaCl, 1 mM EDTA) until the protein concentration in the fractions was below 0.02 mg/ml. Subsequently, Strep-tagged proteins were eluted using 1 ml buffer W containing 2.5 mM desthiobiotin. The elution fractions were evaluated by SDS-PAGE.

The Bio-Rad dye-binding assay was used to determine protein concentrations. Bovine serum albumin was used as standard.

### RNA fishing by protein RNA co-precipitation

To identify initial RNA interaction partners for RnpM and KhpA, the strains GP3753 and GP3752 that harbor genomic deletions of *rnpM* and *khpA*, respectively, were transformed with pGP3617 (Strep-RnpM) and pGP3616 (Strep-KhpA), respectively. For the protein control, strain GP3751 (Δ*ptsH*) was transformed with pGP961 (Strep-PtsH) and for the positive control, GP469 (Δ*csrA*) was transformed with pGP381 (Strep-CsrA). To validate the proposed RnpM binding area of the P RNA, the strain GP3753 was transformed with pGP3686 (Strep-RnpM^K11Y,T15W^) and pPG3697 (Strep-RnpM^V13Y,D53W^). Strains were cultured and expressed as described for Strep-tagged proteins above.

Buffer W without EDTA, pH7.5 (100 mM Tris-HCl, 150 mM NaCl) was used to resuspend and purify the Strep-tagged proteins. Strep-tagged proteins together with potential attached RNAs were eluted by adding one column volume of buffer E (buffer W without EDTA, 2.5 mM d-desthiobiotin) over the columns for three times, respectively. The elution fractions were collected in tubes that were previously supplemented with 2 µl Protector RNase inhibitor (Roche Diagnostics, Karlsruhe, Germany). The elution fractions were split, since 350 µl of each fraction were used for RNA extraction and the remaining protein subjected to SDS PAGE followed by Coomassie staining to evaluate the expression. To extract the RNA from the elution fractions, phenol-chloroform-isoamylalcohol (25:24:1) was added to the elution fractions in a 1:1 ratio. After mixing the samples for 1 min on a vortex device, they were transferred to 2 ml Phase Lock gel heavy tubes (PLG; 5 PRIME), incubated for 2 min and then centrifuged for 30 min at 14,800 rpm and 4°C. The resulting supernatants were transferred into fresh reaction tubes, three volumes of ice-cold 96% EtoH:4M LiCl (30:1) and 1 µl Glycoblue (Invitrogen, USA) were added and mixed by inverting the tubes. Afterwards, the samples were incubated at -20°C overnight to precipitate the RNA. Next, the samples were centrifuged for 30 min at 14,800 and 4°C, the supernatants were discarded, and the remaining pellets were washed two times with ice-cold 70% EtOH. The washed RNA pellets were then air-dried under the fume hood and, subsequently, dissolved in 33 µl RNase-free H_2_O for 1 h at 37°C with agitation. The elution fractions containing the pure target protein, according to the SDS-PAGE, were pooled and digested with DNase I for 2 h at 37°C and 750 rpm. DNase I digestion was monitored by check PCR and followed by a final phenol-chloroform-isoamylalcohol extraction to exclude the DNase I. Finally, the purified RNA was subjected to Ilumina RNA sequencing.

For sequencing, the strand-specific cDNA libraries were constructed with a NEB Next Ultra II Directional RNA library preparation kit for Illumina and the NEB Next Multiplex Oligos for Illumina (96) (New England BioLabs, Germany). To assess quality and size of the libraries, samples were run on an Agilent Bioanalyzer 2100 using an Agilent High Sensitivity DNA Kit as recommended by the manufacturer (Agilent Technologies). Concentration of the libraries was determined using the Qubit® dsDNA HS Assay Kit as recommended by the manufacturer (Life Technologies GmbH, Germany). For sequencing the MiSeq instrument (Illumina) was used in the paired-end mode and 2×75 cycles (v3 chemistry).

The reads obtained from RNA sequencing were processed and analyzed using Geneious Prime software (Geneious Prime 2020.2.4, Biomatters Ltd., New Zealand, 36). Two single read files per sample were merged (paired-end, insert size set to 500) and mapped to the *B. subtilis* 168 genome (37). Enriched RNAs were identified by analyzing the mapped reads using “Find High/Low Coverage” tool (settings: regions with high/low coverage above/below standard deviations from mean: 2).

### Preparation of protein and RNA molecules

The preparation of recombinant His-tagged P protein from *B. subtilis* was done as described for the *E. coli* P protein (38). Tag-free RnpM was prepared from *E. coli* strain Rosetta(DE3) carrying pGP3618 as described above. T7 runoff transcription and 5′-endlabeling was performed as described (39). For transcription of ptRNA^Gly^ as well as *B. subtilis* P RNAs, see (38).

### Pulldown assay to analyse a potential RnpA – RnpM interaction

In order to analyse a potential interaction between RnpA and RnpM, *E. coli* Rosetta (DE3) was transformed with pGP3650 (Strep-RnpM) and the protein was expressed and purified as described above until the washing step was reached. After that, crude extract obtained from *E. coli* Rosetta (DE) harboring the plasmid pQE30-His-RnpA (40) was added to the column. After a subsequent washing step similar to the prior washing, the Strep-RnpM was eluted from the column using buffer W containing 2.5 mM desthiobiotin. The wash and elution fractions were analyzed by a Western blot using antibodies specific for His- and Strep-tag, respectively.

### Bacterial two-hybrid assay

As an alternative approach to identify protein-protein interactions, a bacterial two-hybrid (BACTH) analysis was performed (41). The BACTH approach is based on the interaction-mediated reconstruction of the *Bordetella pertussis* adenylate cyclase (CyaA) activity in *E. coli* BTH101. RnpB and RnpM were fused to the T25 and T18 domain of the adenylate cyclase. An association of the two-hybrid domains leads to functional complementation and thus, cAMP synthesis. This was monitored by detection of the ß-galactosidase activity of the cAMP-CAP-dependent promotor of the *E. coli lac* operon. The plasmids pUT18, pUT18C and p25N, pKT25 allow the expression of proteins fused to the T18 and T25 fragments of CyaA, respectively. The plasmids pGP3672-pGP3675 were prepared for the C- and N-terminal *rnpM* fusions to the T18 and T25 domains, respectively. Similarly, the plasmids pGP3676-pGP3679 were constructed for *rnpA*. The BACTH was initiated by the co-transformation of *E. coli* BTH101 with the prepared plasmids and resulting protein-protein interactions were then analysed by plating the cells on LB plates containing 50 µg/ml kanamycin, 100 µg/ml ampicillin, 40 µg/ml X-Gal (5-bromo-4-chloro-3-indolyl-ß-D-galactopyranoside) and 0.5 mM IPTG (isopropyl-ß-D-thiogalactopyranoside). The plates were incubated at 30°C for a maximum of 48 h.

### RNase P activity assays

RNase P holoenzyme kinetics were determined in buffer KN (20 mM HEPES-KOH, pH 7.4, 2 or 4.5 mM Mg(OAc)_2_, 150 mM NH_4_OAc, 2 mM spermidine, 0.05 mM spermine and 4 mM β-mercaptoethanol) (42). *In vitro* reconstitution of RNase P holoenzyme was performed as follows: 10 nM of P RNA was incubated in KN buffer for 5 min at 55°C and 50 min at 37°C, after which RNase P protein or RnpM protein (each 20 nM) was added in two different assay setups. First, RnpM was added to the previously assembled holoenzyme complex consisting of P RNA and RnpA, alternatively, RnpM was added to *in vitro* reconstituted P RNA prior to addition of RnpA. Finally, the assay sample is incubated for another 5 min at 37°C, before addition of the substrate. The substrate ptRNA^Gly^ was added with a final concentration of 100 mM, 5’-endlabeled substrate was added in trace amounts (<1 nM). Processing reactions were started by combining enzyme and tRNA substrate solutions (final volume 10 µl) and assayed for 0 seconds, 40 seconds, 4 minutes and 12 minutes at 37°C, respectively (40).

### Analysis of RNA-protein interactions

To investigate the interaction between the RNase P RNA with its cognate P protein and RnpM in more detail, we performed *in vitro* cross-linking followed by mass spectrometry to identify the sites in the proteins that bind the P RNA. Protein-RNA complexes were reconstituted as described above and crosslinked with 1 mM mechlorethamine (nitrogen mustard) or 10 mM 1,2,3,4-diepoxybutane for 10 minutes at 37 °C. Unreacted crosslinker was quenched with 13.2 µl 1 M Tris-HCl pH 7.5 and the samples were ethanol precipitated.

The resulting pellets were dissolved in 4 M Urea, 50 mM Tris-HCl pH 7.5 and diluted to 1M with 50 mM Tris-HCl pH 7.5. RNA digestion was performed with 1 µl of Universal Nuclease (Pierce), Nuclease P1 (New England Biolabs), RNAse T1 (Thermo Scientific), RNase A (Thermo Scientific) for 3 hours at 37°C. Protein digestion was achieved with 2.5 µg of trypsin (Promega) and overnight incubation at 37°C. Non-crosslinked nucleotides were depleted with the use of C18 reversed-phase MicroSpin columns (Harvard Apparatus). Bound peptides were eluted with successive application of 50% (v/v) acetonitrile (ACN), 0.1% (v/v) formic acid (FA) and 80% (v/v) ACN, 0.1% (v/v) FA. Enrichment of crosslinked peptides was done with self-packed TiO_2_ spin columns in the presence of 5% (v/v) glycerol as competitor as previously described (43). The eluate was dried under vacuum and resuspended in 15 µl 2% (v/v) ACN, 0.05% (v/v) trifluoroacetic acid, of which 6 µl were used for LC-MS analysis.

Chromatographic separation of crosslinked peptides was done with Dionex Ultimate 3000 UHPLC (Thermo Fischer Scientific) coupled with C18 column packed in-house (ReproSil-Pur 120 C18-AQ, 3 µm particle size, 75 µm inner diameter, 30 cm length, Dr. Maisch GmbH). A 44 minute linear gradient was formed with mobile phase A (0.1% v/v FA) and B (80% ACN, 0.08% FA) from 8% to 45% mobile phase B. Eluting peptides were analyzed with Orbitrap Exploris 480 (Thermo Fischer Scientific). Survey scans were acquired with the following settings: resolution - 120 000; scan range – 350-1600 *m/z*; automatic gain control target – 250%; maximum injection time – 60 ms. Analytes with charge states 2 to 7 were selected for fragmentation with normalized collision energy of 28%. Dynamic exclusion was set to 9s. MS/MS spectra were acquired with the following settings: isolation window -1.6 *m/z*; resolution – 30 000; automatic gain control target – 100%; maximum injection time – 120 ms.

The resulting .raw files were analyzed with the OpenNuXL node of OpenMS (version 3.0.0) (44). Default general settings were used and “RNA-NM Extended” and “RNA-DEB Extended” were selected as a preset. The sequences of the sample protein and commonly observed contaminants supplied with MaxQuant version 2.1.4.0. were provided as a database. Spectra with false discovery rates of 1% and localization scores between 1.0 and 2.0 were selected for molecular modeling.

### Molecular modeling of protein-RNA interaction

An atomic model of *B. subtilis* RnpM has been created with the AlphaFold Monomer v2.0 pipeline (45). The RnpB RNA (RNA component of RNase P) has been modelled using Rosetta’s Fragment Assembly of RNA with Full-Atom Refinement (FARFAR) method integrating RNA-Puzzle-inspired innovations (FARFAR2) (46). Three crystal structures have been used as templates: 417-nt ribonuclease P RNA from *Bacillus stearothermophilus* (PDB id: 2A64, 80% sequence identity, region 11-403, 47), the specificity domain of RNase P RNA of *B. subtilis* (1NBS, 100% sequence identity, region 80-239, 48) and the RNase P holoenzyme of *Thermotoga maritima* (3Q1R, loops 248-255, 333-354, 49). The resulting atomic model of RnpB has been improved in terms of the geometrical quality by the ERRASER-Phenix pipeline using a physically realistic model of atomic-level RNA interactions (50). This pipeline was used to resolve potential steric clashes and anomalous backbone and bond geometries of the RnpB RNA model that could affect the scoring function employed for assessment of subsequently performed *in silico* docking experiments. The crystal structure of the *B. subtilis* RNase P protein component (1AF6, 51) was edited by removal of solvent components and used to assemble a complex with the *B. subtilis* P RNA model based on superposition with corresponding compartments of the *T. maritima* RNase P holoenzyme in complex with tRNA (3Q1Q, 49). Blind docking of RnpM to P RNA was conducted with ROSETTA tools designed to model structures of RNA-protein complexes (fold-and-dock method) and the stand-alone version of HDOCKlite (52, 53). The resulting models were analysed using self-written tools to assess their compatibility with spatial restraints derived from crosslink data obtained using the chemical crosslinking agents 1,2,3,4-diepoxy-butane (DEB) and nitrogen mustard (NM). These tools supported a multi-core environment when executed using GNU parallel software (https://www.gnu.org/software/parallel/). Individual mass spectrometry experimental datasets, obtained using several localization scores (LS: 1.0, 1.5, 2.0), were filtered to comprise only those crosslinks which have been observed at least two times (redundancy ≥ 2).

Almost two hundred thousand docking models have been checked for their individual compatibility with each of the three sets of spatial restraints derived from mass spectrometry datasets (LS: 1.0, 1.5, 2.0). It has been achieved by calculating Euclidean distances between the protein residue (position within the protein is known) and a directly linked nucleotide/nucleotides (position within the tRNA is not known) that is/are located within the specified distance from the protein residue. The applied distance thresholds were dependent on the cross-linker type (15.0 Å for DEB, 10 Å for NM), and have been measured between the atom N3 (nucleotides U or C) or N7 (remaining nucleotides) of RNase P RNA compartment and Cα position of the cross-linked protein residue. In that manner both single (d_i_0, the shortest-distance, one per protein residue) and multiple crosslinker occurrences (d_i_0-ALL) between the particular protein residue and all potentially crosslinked nucleotides have been assessed, counted and stored. To account for flexibility of protein and RNA molecules, slacked distance thresholds (2 Å, 3 Å, 5 Å) were additionally applied leading to assessment of d_i_2-ALL, d_i_3-ALL and d_i_5-ALL, respectively. These “per protein residue” assessed number of potentially crosslinked nucleotides (d_i_X, d_i_X-ALL; X=2,3,5) have been summed up for each particular model (decoy) leading to the overall number of crosslinks per modeled RnpM-RNase P complex (d0, d2, d3, d5, d0-ALL, d2-ALL, d3-ALL, d5-ALL, see Supplementary Table S3). The latter were used to find models supporting the highest number of potential crosslinks. In addition, all decoys have been characterized using a simple assessment of RnpM-RNase P assembly quality that was based on three attributes calculated between components of the modeled complex: the buried surface area (54), number of hydrogen bonds and unfavorable interatomic distances (clashes, shorter than 2.2 Å) (contact program from the CCP4 suite) (55). The last attribute (the number of clashes ≤ 10) was used to filter out decoys that could reveal artificially high number of crosslinks due to non-physical assembly of the RnpM and RNase P compartments. The filtered list of decoys has been subjected to a sorting routine that served to the identification of decoys belonging to the top scoring models in the following distance-based categories (d0, d0-ALL, d1, d2-ALL, d3, d3-ALL, Supplementary Table S4). The remaining number of potential crosslinks (d5 and d5-ALL), were left aside for potential validation purpose. Taken together, the sorting procedure has been performed for three MS crosslink data sets obtained using minimal localization scores (LS) of 1.0, 1.5, and 2.0 and included decoys with the number of intermolecular clashes ≤ 10.

## RESULTS

### Identification of the RNase P RNA as binding partner of RnpM (YlxR)

A recent global study on RNA-binding proteins in the Gram-positive pathogen *C. difficile* identified paralogs of the *B. subtilis* proteins KhpA and RnpM (YlxR) as novel RNA-binding proteins (11). To study whether these proteins interact with RNA in *B. subtilis* and to identify potential interacting RNAs, we decided to purify them together with their bound RNA molecules and to identify associated RNA molecules by RNA-seq. For this purpose, we fused the KhpA and RnpM proteins to a C-terminal Strep-tag. As controls, we used the PtsH protein that is known to interact with many other proteins but not with RNA, as well as CsrA that binds specifically to the *hag* mRNA in *B. subtilis* (56), similarly fused to a Strep-tag. To avoid interference of the native non-tagged proteins, the plasmids for the expression of the fusion proteins were propagated in the corresponding gene deletion mutants. The results are shown in Fig. 1. An analysis of the RNA interactome of PtsH yielded only the highly abundant ribosomal RNAs which account for about 85% of the total cellular RNA. These ribosomal RNAs were retained at similar levels in all analyses, irrespective of the bait protein, suggesting non-specific binding. For CsrA, we identified a single specifically interacting RNA, the *hag* mRNA. This is in excellent agreement with previous results (56). These results indicated that our experimental system is well suited to identify RNA molecules that bind specifically to the bait proteins of interest. For KhpA we did not co-purify any specific RNA molecules indicating that this protein either (i) does not bind RNA in *B. subtilis*, (ii) that it interacts with RNA molecules under rather specific conditions, or (iii) that it binds RNA species of low abundance that escaped detection. For RnpM, we identified a single RNA species in addition to the non-specifically binding rRNAs, the RNA subunit of RNase P, also called RnpB. Based on the results obtained with the positive and negative controls, we considered the RnpM protein as a specific interaction partner of RnpB RNA. The interaction might, however, be mediated by the RNase P protein component, RnpA.

**Figure 1.**
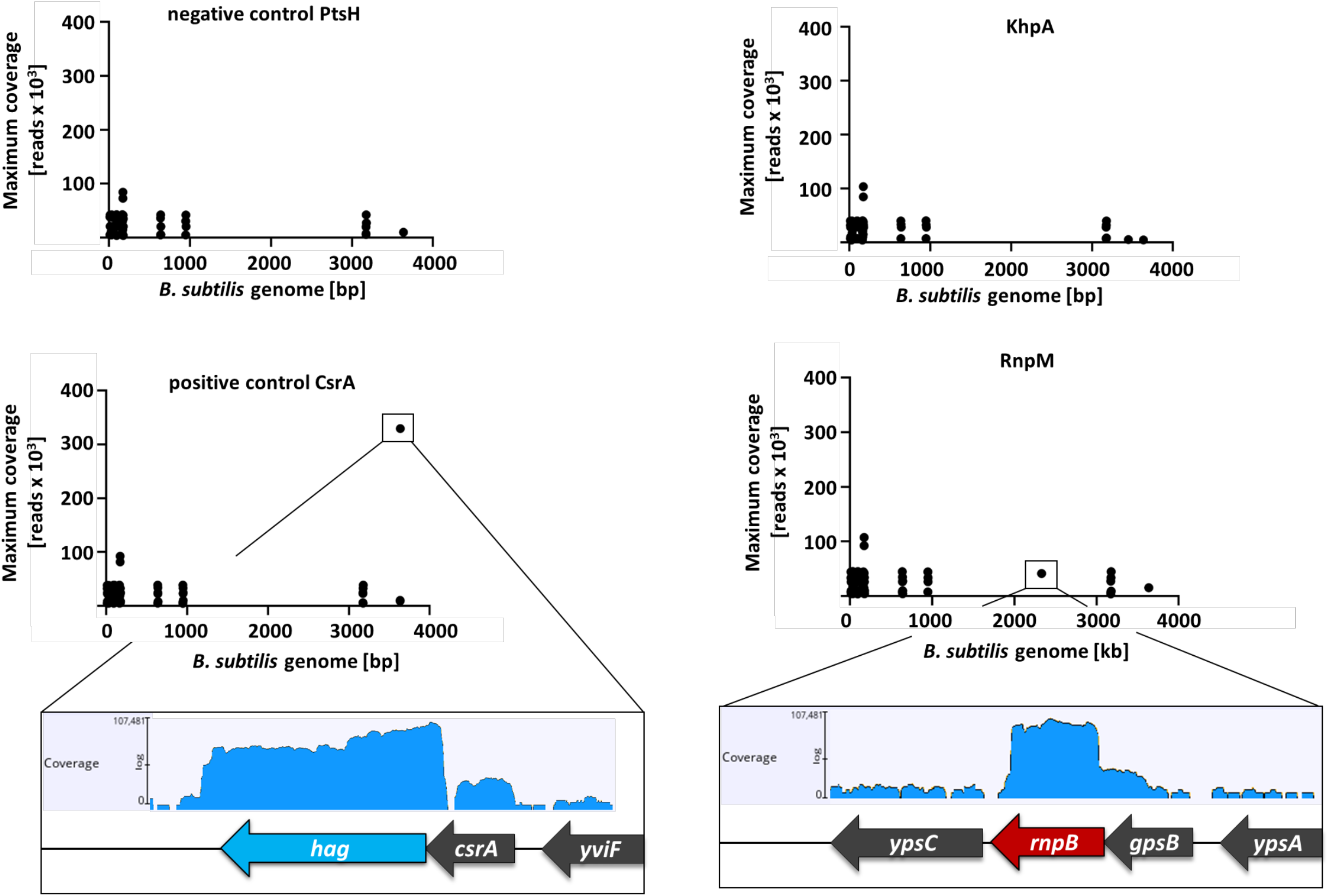
RnpM interacts with RNase P RNA. Protein-RNA co-precipitation approach with GP3751 (Δ*ptsH*) + pGP961 (Strep-PtsH), GP469 (Δ*csrA*) + pGP381 (Strep-CsrA), GP3752 (Δ*khpA*) + pGP3616 (Strep-KhpA), and GP3753 (Δ*rnpM*) + pGP3617 (Strep-RnpM). Strep-tagged proteins were expressed and purified via a Strep-Tactin matrix and RNA extracted from the elution fractions, respectively. Enriched RNAs were identified by mapping the reads obtained from RNA-sequencing onto the *B. subtilis* genome.

### RnpM finetunes the activity of RNase P

RNase P catalyzes the removal of 5’-leader sequences from precursor tRNAs to generate the mature tRNAs. The enzyme consists of the catalytically active RNA (RnpB) and the structurally conserved protein subunit (RnpA) that is important for substrate recognition (22). The interaction of RnpM with the RNase P RNA component suggested that RnpM might modulate the activity of RNase P. To address this question, we performed RNase P activity assays using purified components of RNase P, RnpM, and a bacterial precursor-tRNA^Gly^ (pre-tRNA^Gly^) with a 14-nt long 5’ leader as the substrate. We purified His-RnpA and tag-free RnpM, reconstituted the holoenzyme with *in vitro*-transcribed RnpB RNA, and assayed the processing of radioactively labelled 5’-P^32^-pre-tRNA^Gly^ (see Materials and Methods for details) in the presence or absence of RnpM. The assay was performed in two distinct setups: in the first setup, we reconstituted RNase P by preincubating the RNA with RnpA protein, followed by addition of substrate alone or substrate and RnpM (see Fig. 2A). In the second setup, we first pre-incubated the RNA with RnpM before addition of RnpA to allow RnpB-RnpM complex formation that may interfere with holoenzyme assembly (see Fig. 2B). For each setup, five independent experiments were performed to determine average processing rates (mean *k*_obs_ values). In both scenarios, the addition of RnpM resulted in a reduction of RNase P activity by about 25 to 33%. The difference was most pronounced and statistically significant when the RnpB RNA was pre-incubated with RnpM prior to the addition of the RNase P protein. These results indicate that RnpM is able to finetune RNase P activity.

**Figure 2.**
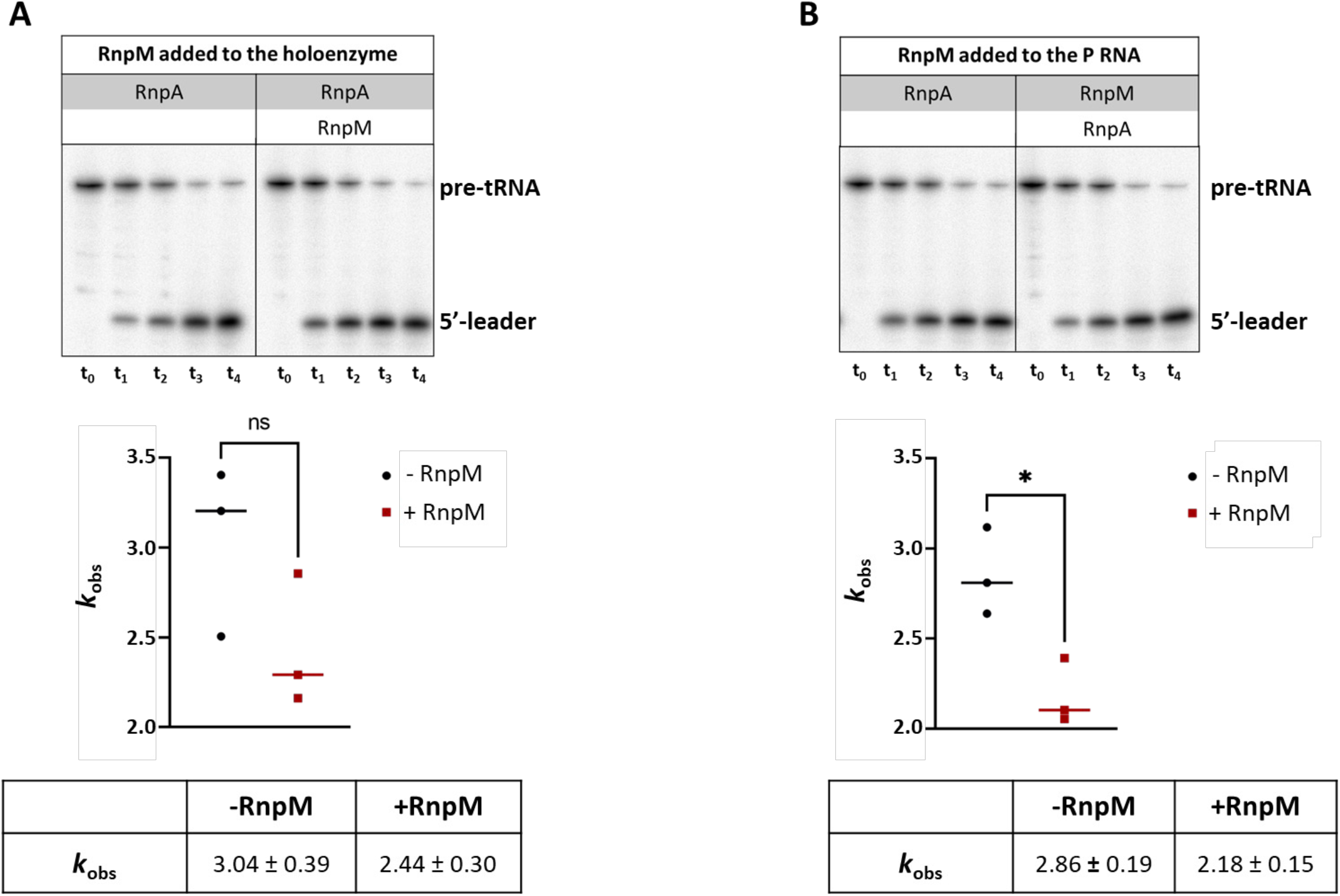
RnpM negatively affects RNase P activity. Cleavage activity of RNase P acting on ptRNA^Gly^ was assayed *in vitro* in presence and absence of RnpM protein, respectively. RNase P RNA was used in a final concentration of 10 nM, the proteins RnpA and RnpM with a final concentration of 20 nM. Assay conditions: 20 mM HEPES pH 7.4, 4.5 mM Mg(OAc)_2_, 150 mM NH_4_OAc, 2 mM spermidine, 0.05 mM spermine, 4 mM β-mercaptoethanol. The substrate ptRNA^Gly^ was added with a final concentration of 100 mM, 5’-endlabeled substrate was added in trace amounts (<1 nM). K_obs_ represents pmol substrate converted per pmol of P RNA per min. The effect of RnpM on RNase P cleavage activity was assessed in two setups: (A) RnpM was added to the pre-reconstituted RNase P holoenzyme. (B) RnpM was added to P RNA prior to addition of the P protein RnpA.

### RnpA and RnpM do not interact with each other

The results described above demonstrate (i) that the *B. subtilis* RnpB RNA and the RnpM protein physically interact with each other, and (ii) that RnpM is a moderately negative effector of RNase P activity. However, these results did not allow a final conclusion about the mechanism of this modulation. The interaction of RnpM with RNase P RNA might be mediated by RnpA, the protein subunit of RNase P, the two proteins might compete for binding to the RnpB RNA, or binding of RnpM might interfere with the catalytic activity of RNase P.

The RnpA and RnpM proteins share several features, among them the ability to bind the P RNA, their surface charge distribution, and their expression pattern. Both proteins possess large positively charged surfaces and a positively charged cleft that might interact with nucleic acids (see Fig. 3 A). Typically, protein-protein interactions are meaningful if the cellular amounts of the partners are in a certain range. Based on an analysis of protein quantities in *B. subtilis* (Reuss 2017), the amount of RnpM is about 3.5 times higher than that of RnpA. This would be compatible with a role for RnpM in the control of RnpA binding activity. We also compared the expression of the *rnpA* and *rnpM* genes under more than 100 conditions (see Fig. 3 B). Under all conditions, *rnpM* is more strongly expressed than *rnpA*. Interestingly, the expression levels of the two genes show a very strong positive correlation (4), again indicating a functional link.

**Figure 3.**
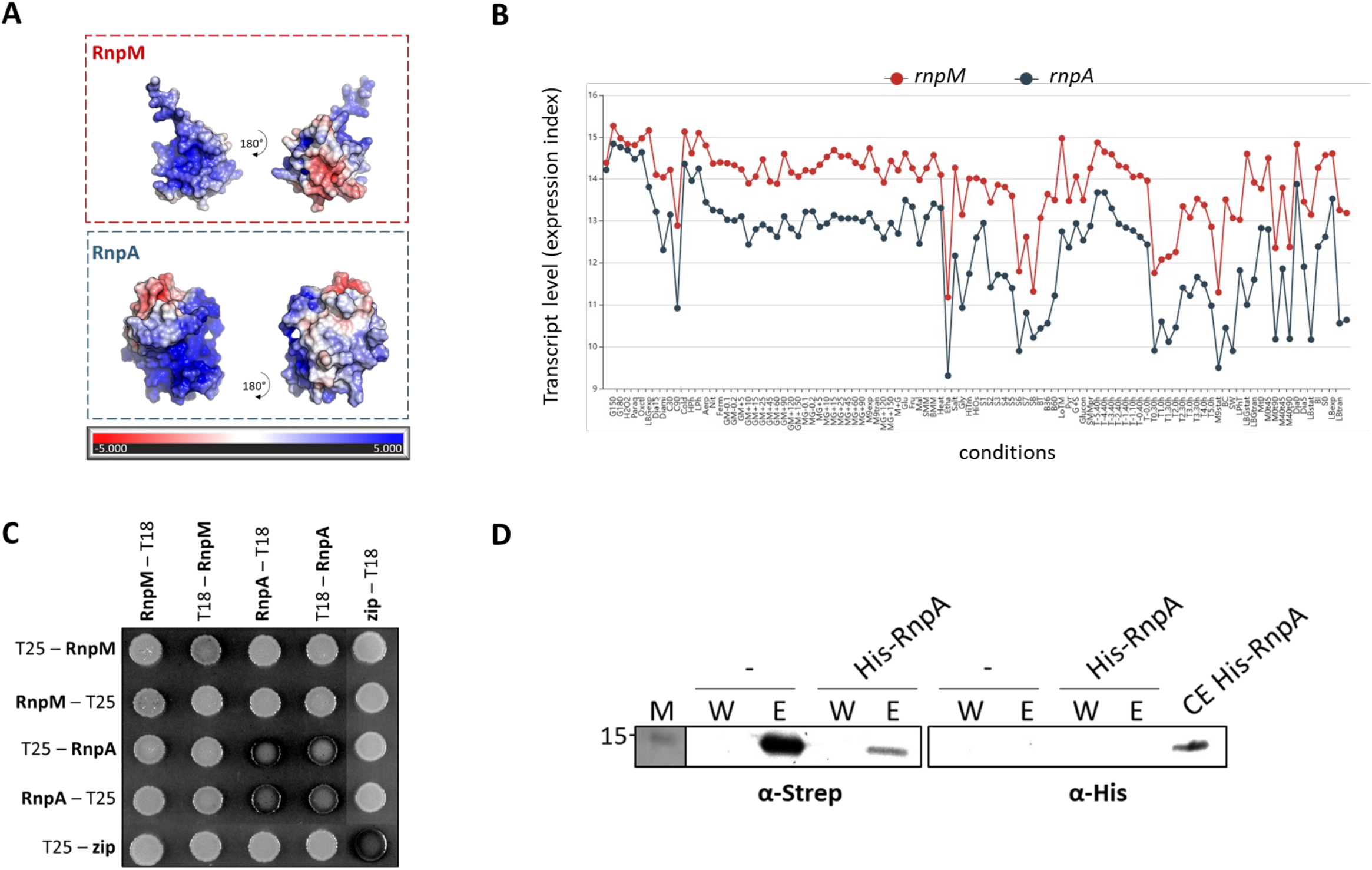
Comparative analysis of RnpM and RnpA. (A) Electrostatic surface charge distribution was analyzed for RnpM (AF-P32728-F1) and RnpA (PDB: 1A6F) using APBS software (74). The surface potential is ramp-colored from red to blue (-5 to 5 kT/e). (B) Expression of *rnpM* (red) and *rnpA* (blue) was assessed by comparing the transcript levels of the two genes under more than 100 conditions (4). For more detailed information on the tested conditions, see http://www.subtiwiki.uni-goettingen.de/v4/expression?gene=F4097349A563503468A2A14F062AEAC532C7917A. (C) To test for a potential physical interaction between RnpM and RnpA, a bacterial two-hybrid (BACTH) experiment was performed, dark colonies indicate an interaction. (D) An additional *in vitro* pulldown approach was performed using immobilized and purified Strep-tagged RnpM immobilized on a Strep-Tactin column and passing buffer W (-) or *E. coli* Rosetta cell extract expressing the *B. subtilis* His-RnpA (His-RnpA) through the column. The eluates (E) and wash fractions (W) were analyzed by Western blot analysis using antibodies against the Strep- and the His-tag, respectively. The position of a 15 kDa marker protein loaded on such SDS gels is indicated on the left.

To address the potential functional link of RnpM and RnpA, we tested the possibility of a protein-protein interaction between the two proteins. The potential physical interaction between RnpM and RnpA was first tested using a bacterial two-hybrid system that is based on the reconstitution of adenylate cyclase and the resulting expression of the lac operon in *E. coli* upon interaction of the two partner proteins. RnpA showed a strong self-interaction, as expected based on previous results obtained with the protein of *Staphylococcus aureus* (57). In contrast, no interaction between RnpA and RnpM was detectable (Fig. 3 C). In a second approach, we expressed and purified Strep-tagged RnpM, immobilized the protein to a Strep-Tactin column and applied an *E. coli* cell extract expressing the recombinant His-tagged RnpA from *B. subtilis*. The Strep-RnpM protein was then eluted by the addition of desthiobiotin, and the elution fraction was tested for the presence of RnpA. As shown in Fig. 3 D, the Strep-tagged RnpM protein was detected in the elution fractions, if the column was incubated with buffer W or with the cell extract prior to elution. In contrast, His-RnpA did not co-elute with RnpM even though the protein could be detected in the cell extract (Fig. 3D, lane CE His-RnpA). These results suggest that the two proteins do not physically interact with each other. However, interaction screens are typically prone to false-negative results (58). Therefore, we also considered a recent proteome-wide protein-protein interaction study with *B. subtilis* based on *in vivo* cross-linking and complex enrichment. Even though both proteins are highly abundant in *B. subtilis* (4), they were not involved in any protein-protein interaction (59).

Thus, our results suggest that RnpM does not interact with the RnpA protein. The observed modulation of RNase P activity by RnpM is therefore likely to be achieved in a RnpA-independent manner.

### Chemical cross-linking of RnpB and RnpM suggests areas of molecular interaction

To test whether RnpM directly binds to RNase P RNA, we incubated *in vitro* transcribed P RNA with purified RnpM, chemically cross-linked potential complexes using 1,2,3,4-diepoxy-butane (DEB) and nitrogen mustard (NM) and determined the amino acids of RnpM that interact with the RNA. As a control, we performed the same experiment for the reconstituted RNase P holoenzyme. As expected, both RnpA and RnpM yielded a large number of cross-links thus confirming the physical interaction between the P RNA and either protein (see Supplementary Tables S1, S2).

The cross-linking data were used to model two complexes, the RNase P holoenzyme and the RnpM-P RNA complex. In case of the holoenzyme, the model based on the superposition of *T. maritima* RNase P (PDB 3Q1R, 49) supported the experimentally observed cross-links, hence molecular docking was not required (see Fig. 4). For the RnpM-P RNA complex, *in silico* blind docking was performed. The resulting decoys were checked for compatibility with three cross-linked data sets obtained using localization scores 1.0, 1.5, and 2.0. The analysis revealed that the cross-link data set obtained with LS 1.0 does not yield convergent results (see Supplementary Fig. S1A), however the top-populated cluster of seven RnpM molecules is located close to the RNase P protein and tRNA binding sites (see Fig. 5, Supplementary Fig. S1B). These seven decoys form the group of models with the largest buried surface area, the greatest number of hydrogen bonds and the lowest number of clashes implicating potential biological relevance (see Supplementary Table S4). Interestingly, mass spectrometry data sets obtained with LS 1.5 and 2.0 (see Supplementary Fig. S1C and D) reveal only three potential positions of RnpM on the P RNA; however, binding of RnpM to the RNase P specificity domain or to the area that partially overlaps with the binding site of the RNase P protein are rather unlikely. The first is very distant (on the other side of the P RNA) to the position where tRNA should bind, hence direct interactions or short-range allosteric interactions could be excluded. The latter would indicate a competing binding mechanism with the RNase P protein. The third modeled position of the RnpM molecule overlaps perfectly with the top scoring cluster obtained from the LS 1.0 data set. The predicted RnpM binding site is located close to the tRNA and RNase P protein binding sites (see Fig. 5). This interaction might interfere with productive binding of the pre-tRNA substrate to RNase P.

**Figure 4.**
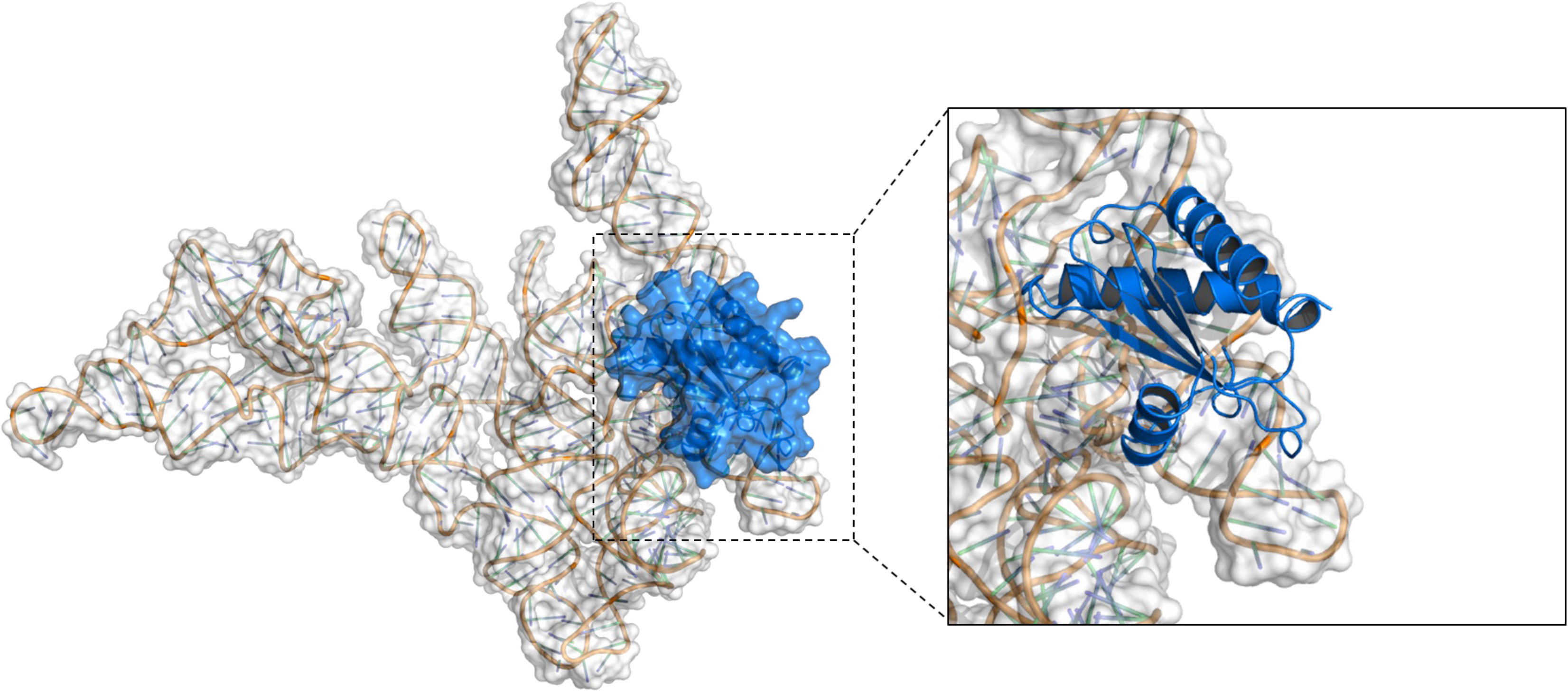
Model of the *B. subtilis* RNase P holoenzyme. The P RNA is depicted as cartoon with a white transparent surface and the P protein is represented by a cartoon in marine blue with a transparent surface. The model is based on superimposition with the *T. maritima* RNase P holoenzyme (PDB: 3Q1R, 49).

**Figure 5.**
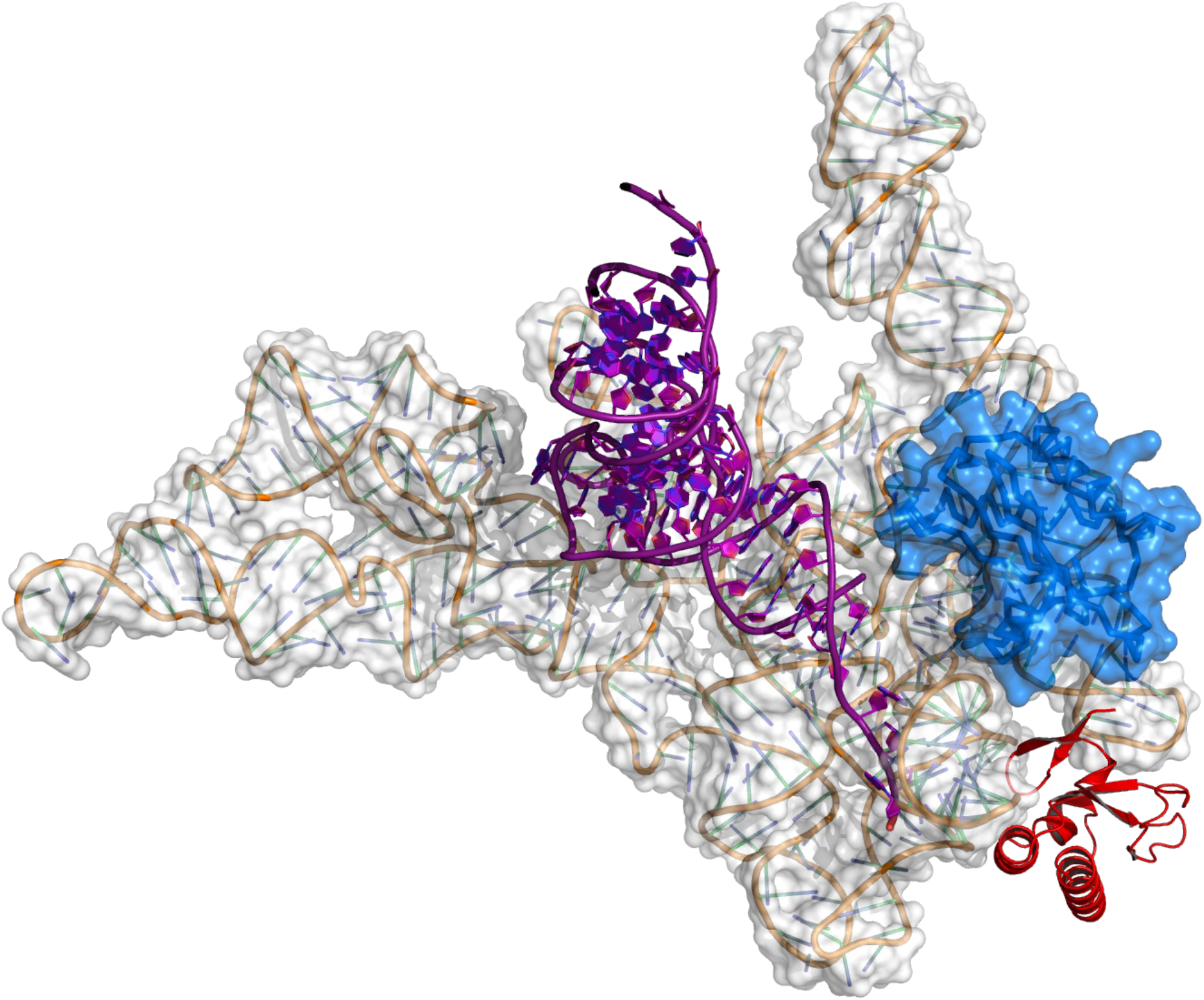
Predicted RNase P-RnpM model in complex with pre-tRNA. The tRNA^Phe^ molecule (purple) has been modeled based on superposition with the crystal structure of the RNase P holoenzyme from *T. maritima* in complex with the tRNA and in the presence of the soaked-in 5’ leader (PDB 3Q1R). The modeled position of RnpM is depicted as a red ribbon diagram.

*In silico* docking yielded a model of the RnpM-RNase P complex supporting spatial restraints derived from cross-linking data. The predicted RnpM binding site is located in the cleft formed by the nucleotides 31-35 of helix P3, nucleotides 245-257 of P15 and nucleotides 271-273 and 285-287 of P15.1 (see (60) for a description of P RNA structural elements). The apical loop of helix P3 occupies the prominent positively charged cleft of RnpM (Fig. 6A) while the other two P RNA elements form interactions with positively charged and neutral patches located on the opposite side of the RnpM molecule (Fig. 6B). The negatively charged patch on the RnpM surface is exposed to the solvent and does not form any interactions with RNase P RNA. Interestingly, the most likely disordered N-terminus of RnpM (residues 1-7), that was not included during *in silico* docking experiments, could potentially form electrostatic and polar interactions with both the RNA and protein subunit of RNase P. However, both the existence and importance of these interactions for binding remain currently speculative. It should be mentioned that docking experiments yielded several hits of the RnpM molecule in the predicted binding site (see Supplementary Fig S2). They exhibit small positional alterations that lead to differences in the number of estimated hydrogen bonds, short molecular contacts and buried surface areas between the interacting molecules (see Supplementary Table S4). The observed heterogeneity of the RnpM position might indicate conformational alterations of RNase P RNA upon binding that in turn might cause changes in the catalytic activity of RNase P.

**Figure 6.**
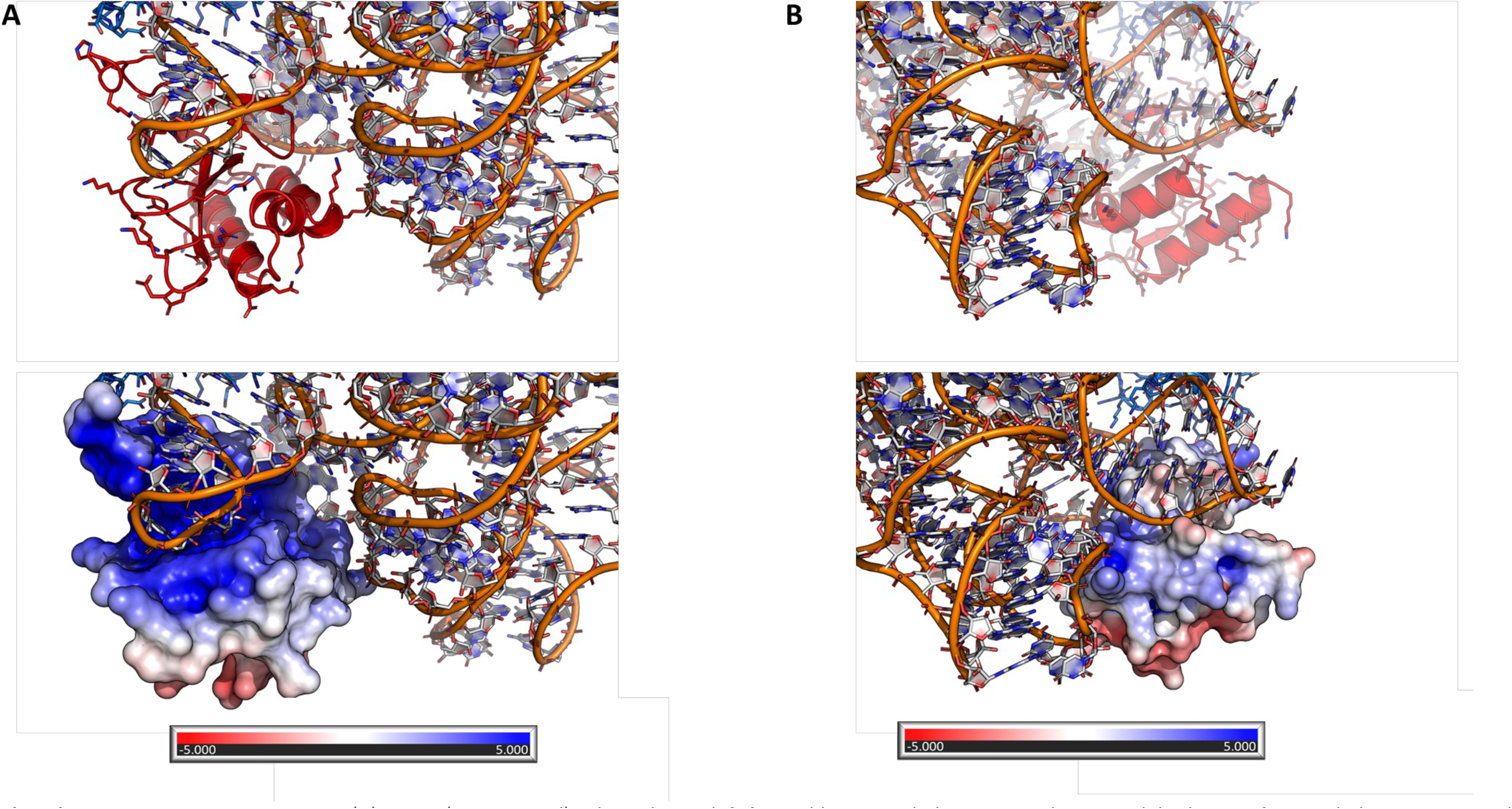
Predicted RnpM-P RNA interaction site. (A) RnpM (cartoon, red) is bound in a cleft formed by P RNA helices P15 and P15.1, while the tip of P RNA helix P3 occupies the positively charged cleft of RnpM. (B) The other two P RNA secondary structures bind to the neutral and positively charged RnpM patches on the other site of the protein.

### Validation of the interaction area

The inherent ambiguity of the docking results was resolved by the introduction of mutations into the RnpM protein and the analysis of their effect on *in vivo* RNA binding. Specifically, we replaced Lys-11 by a tyrosine residue and Thr-15 by a tryptophan (RnpM-1). In a second RnpM variant (RnpM-2), Val-13 and Asp-53 were replaced by tyrosine and tryptophan, respectively. Atomic models of both variants have been predicted using AlphaFold2 and were virtually identical with each other and the wild type RnpM as assessed based on conformation of the main chain atoms (see Fig. 7). Based on our model, we expected no binding to P RNA for the second variant RnpM-2 if the interaction would take place as predicted close to the site of interaction between P RNA and the tRNA substrate. The RnpM-1 variant should still be able to bind to this predicted interaction site (see Fig. 5), although much weaker. Both RnpM variants should not affect interactions with RNase P RNA at the predicted alternative location that would interfere with binding of the P protein, hence if RnpM occupied this latter location, no difference in biochemical assays between wild type and variants should be observed (see Fig. 5, Fig. 7).

**Figure 7.**
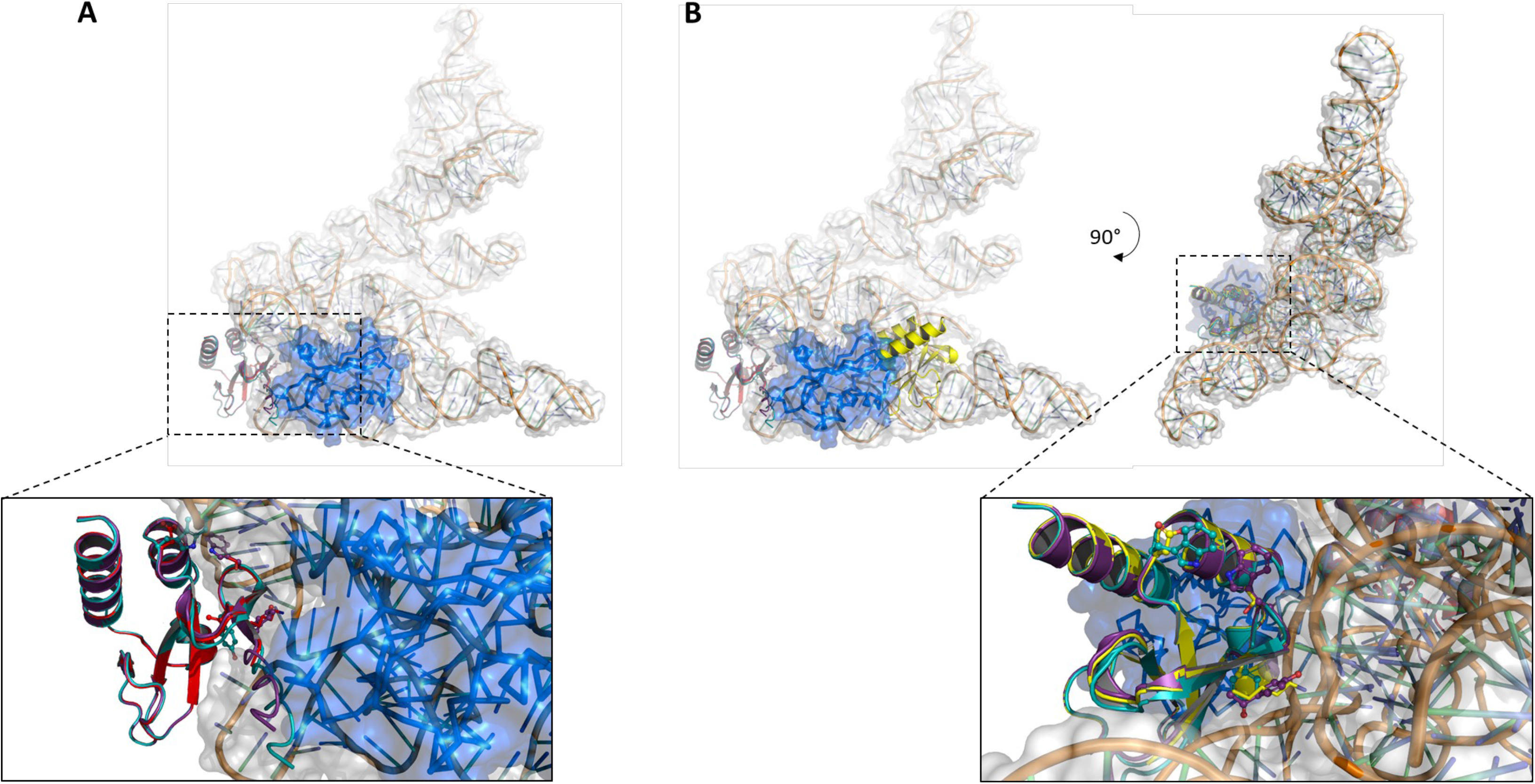
Prediction of atomic models of RnpM variants. Overlay of the wild type protein (red), and the RnpM-1 (purple) and RnpM-2 (cyan) variants. In the RnpM-1 mutant, Lys-11 and Thr-15 were replaced by a tyrosine and tryptophan, respectively. In the RnpM-2 variant, Val-13 and Asp53 were replaced by tyrosine and tryptophan, respectively. (A) The predicted RnpM binding site from Fig. 5, 6 is depicted and (B) an alternative predicted binding site (see Fig. S1) is represented.

The plasmid-borne Strep-tagged mutant proteins as well as the tagged wild type protein were expressed in the *B. subtilis rnpM* deletion mutant, the proteins were purified using a Strep-Tactin column, and RNA molecules bound to the proteins were identified. As observed in our initial experiments (see Fig. 1), all three forms of RnpM showed interactions with ribosomal RNA. This indicates that the mutant variants are correctly folded, as also predicted by AlphaFold2, and able to bind RNA. As shown in Fig. 8, the RnpM-1 protein was still able to bind to the RNase P RNA molecule; however, the enrichment was severely reduced as compared to the wild type protein. In contrast, no P RNA was co-eluted with the RnpM-2 protein. These results are in agreement with the proposed model for the P RNA-RnpM interaction. In contrast, they are not consistent with binding of RnpM to the aforementioned alternative site on P RNA that is also bound by RnpA. Thus, RnpM binds to a cleft of P RNA that is located directly beneath the RnpA protein binding site and that is formed by the structural elements P3, P15 and P15.1. P15 binds the 3’-CCA end of tRNAs (60), hence the identified binding site of RnpM seems to be perfectly positioned for modulating RNase P activity.

**Figure 8.**
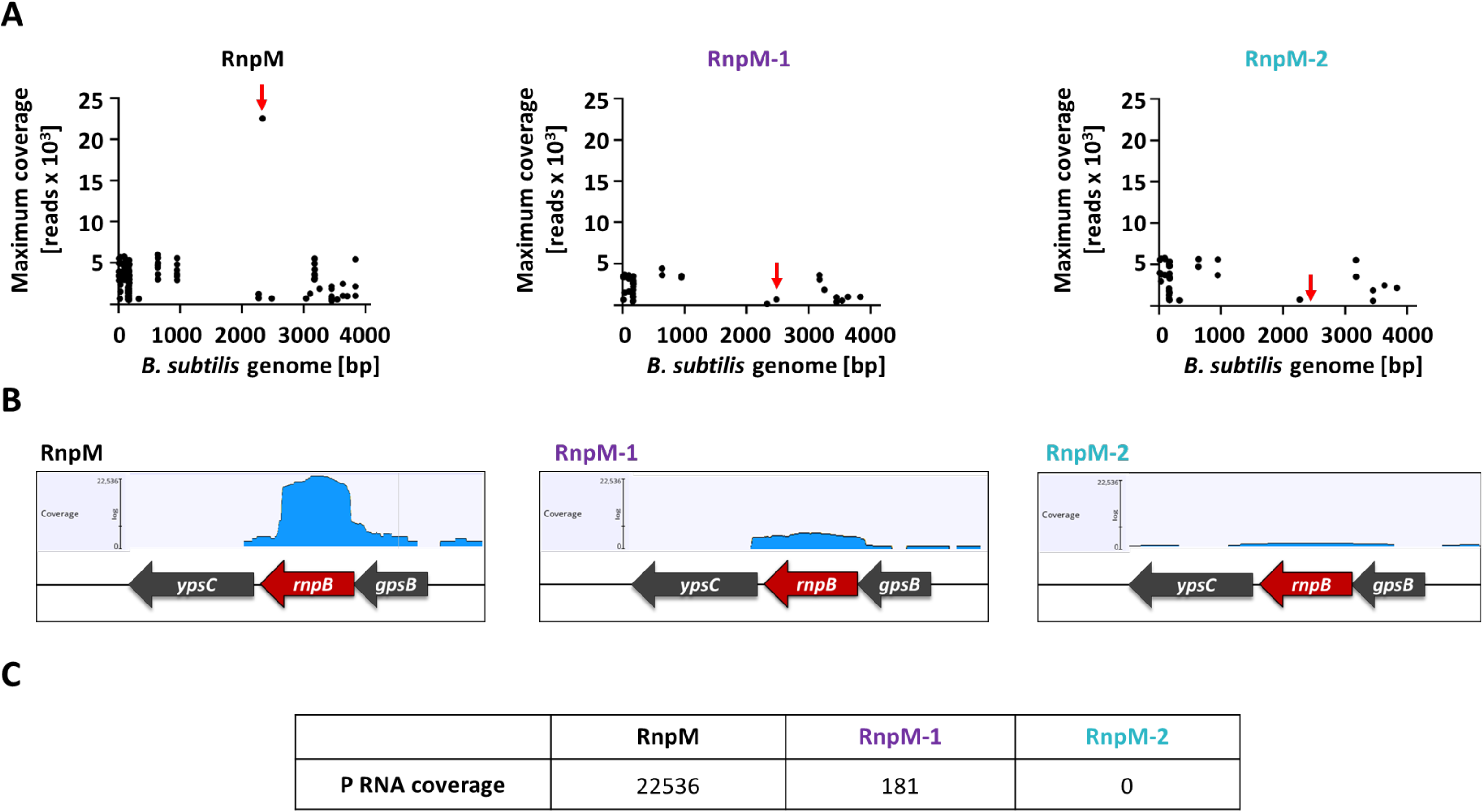
RnpM site directed mutagenesis prevents P RNA binding. (A) Strep-tagged fusion proteins of RnpM, RnpM^K11Y,T15W^ (RnpM-1) and RnpM^V13Y,D53W^ (RnpM-2) were expressed and purified and RNA extracted from the eluates, respectively. Reads obtained from RNA-sequencing were mapped onto the *B. subtilis* genome to identify enriched RNAs. (B) The P RNA binding ability of the RnpM variants was analyzed. (C) Comparison of the P RNA coverage obtained for RnpM, RnpM-1 and RnpM-2 after mapping the reads and specifically screening for P RNA.

## DISCUSSION

The results presented in this study demonstrate that the so far unknown RnpM protein of *B. subtilis* binds to the RNA subunit of the ribonucleoprotein complex RNase P, and that the interaction limits RNase P activity. An analysis of the molecular details of the interaction revealed that RnpM binds close to the area of the P RNA that is required for binding of pre-tRNAs.

RnpM is one of about 40 highly expressed proteins in *B. subtilis*, whose function has so far not been uncovered (5). It has recently been postulated that the identification of functions of unknown proteins requires at least a minimal amount of prior annotation (61). In the case of RnpM, the conserved genomic arrangement with genes involved in transcription and translation, two transcriptome analyses, the crystal structure of the ortholog from *S. pneumoniae*, and the observation that the protein interacts with RNA in *C. difficile* were available prior to this study (11, 19, 20, 21). Nevertheless, no clear function of the protein could be deduced from this knowledge.

Three observations suggest a role of the *B. subtilis* RnpM protein in the cellular information management. First, the gene is part of an operon with several other genes involved in transcription and translation. This operon organization is conserved not only in the Firmicutes, but also in Actinobacteria and Cyanobacteria, indicating a common functional context of the encoded proteins. Second, the corresponding protein from *C. difficile* binds RNA, in good agreement with predictions from the structural analysis (11, 19). However, transcriptome analyses of wild type and mutant strains gave no clear picture and suggested a role for the protein in the expression of a highly diverse set of seemingly unrelated genes. Such a diffuse pattern of changes in gene expression is typical for mutants affected in proteins that are involved in central processes of gene expression such as transcription, translation or the control of DNA topology, and is thus the third hint at a function of RnpM in central information processing. In *B. subtilis*, this has been observed for proteins involved in transcription such as the RNA polymerase β’ subunit or termination factor Rho and for RNases such as polynucleotide phosphorylase, RNase J1 and RNase Y (10, 62, 63, 64, 65, 66). Similarly, highly pleiotropic effects on global gene expression were also found for strains altered in the RNA helicase CshA or the DNA topoisomerase TopA (9,67).

The RnpM protein is nearly ubiquitously present in Gram-positive bacteria as well as in Cyanobacteria and the Alpha-, Delta-, and Epsilonproteobacteria, but it is lacking in the Gammaproteobacteria including *E. coli*, as well as in the Bacteroidetes, and the Spirochaetes. In contrast, two major RNA-binding proteins in the Gammaproteobacteria, Hfq and CsrA, have completely different functions in the Bacilli (CsrA), or the function has not yet been elucidated in the latter group of bacteria (15, 17, 56, 68). Moreover, the functional equivalent of Hfq that acts as the matchmaker for RNA-RNA pairing has not yet been identified in the Bacilli. Thus, similar functions can obviously be taken over by completely different unrelated proteins in different groups of bacteria as exemplified by the unrelated endoribonucleases E and Y which initiate RNA degradation in *E. coli* and *B. subtilis*, respectively (69, 70, 71).

Our work identified the RnpM protein as a modulator of RNase P activity. More specifically, the interaction between the RNase P RNA and RnpM limits the activity of RNase P by interfering with productive pre-tRNA substrate binding. This raises the question why the activity of an essential enzyme that has a key function in the cell, the maturation of tRNAs, has to be controlled. The tRNA genes are typically clustered with the genes for the ribosomal RNAs, and there are multiple genes for individual tRNAs. Thus, pre-tRNAs are produced in high abundance. The limitation of RNase P activity by RnpM suggests an incomplete pre-tRNA maturation. Indeed, the presence of both precursor and mature forms of tRNA has been observed in *E. coli* (72). This may leave reserves in the system that can be used if amino acids and carbon backbones are available in high abundance and when they can be completely allocated to translation because there would be no need for amino acids in catabolic pathways. RnpM is the first example for a protein that modulates RNase P activity in bacteria. It is tempting to speculate that additional mechanisms to control the activity of this important ribozyme will be discovered, particularly in those bacteria that do not possess RnpM.

As mentioned above, research tends to focus on those proteins that have already been functionally characterized. More and more information is published on those proteins. In contrast, there is a gap concerning the investigation of proteins of unknown function. Only the availability of some initial information allows the functional characterization of unknown proteins (61). The RnpM protein is an excellent example that this prediction holds true. With the initial knowledge that RnpM is an RNA-binding protein, experiments could be designed to identify its RNA interaction partner and then the molecular details of the interaction. Given the large number of unknown proteins in any genome, we trust that our experimental pipeline will be fruitful also for the identification of other proteins.

## SUPPLEMENTARY DATA

Supplementary Data are available at NAR Online.

## ACKNOWLEDGEMENTS

We are grateful to Bastian Dörrbecker and Fabian Commichau for their help with plasmid construction. We acknowledge Sabine Lentes and Monika Raabe for technical assistence.

