## Supplementary material for "The previously uncharacterized RnpM (YlxR) protein modulates the activity of ribonuclease P in *Bacillus subtilis*": Fig. S1

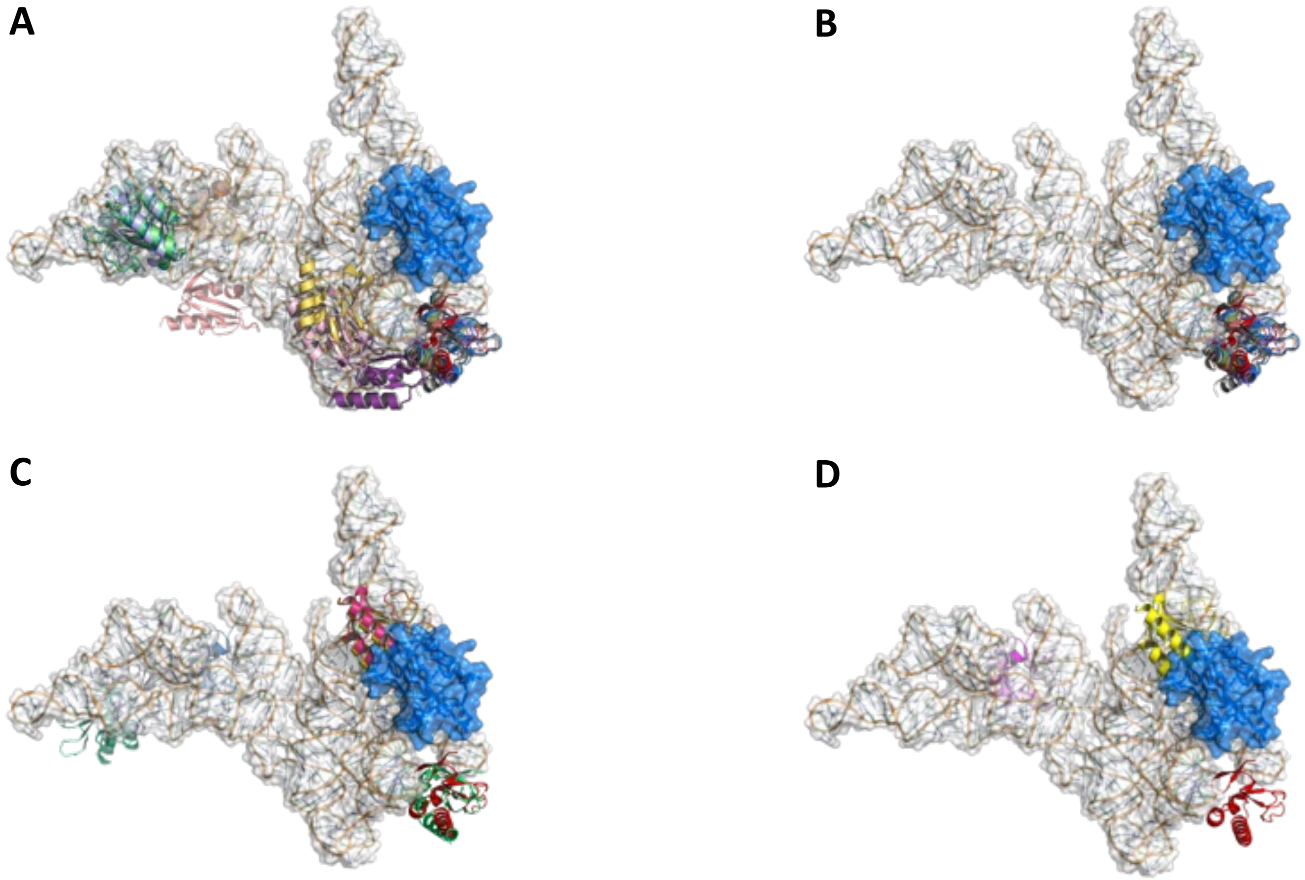

Fig S1. Decoys obtained from blind docking using RnpM-P RNA cross-linking data sets. P RNA and P protein are depicted in transparent white and marine, respectively, and the decoys each in a different color. The decoys obtained from *in silico* blind docking were checked for compatibility with cross-linking data sets with the localization scores (LS) 1.0 (A), 1.0 top populated cluster (B), 1.5 (C) and 2.0 (D).

Wicke et al., Fig. S1

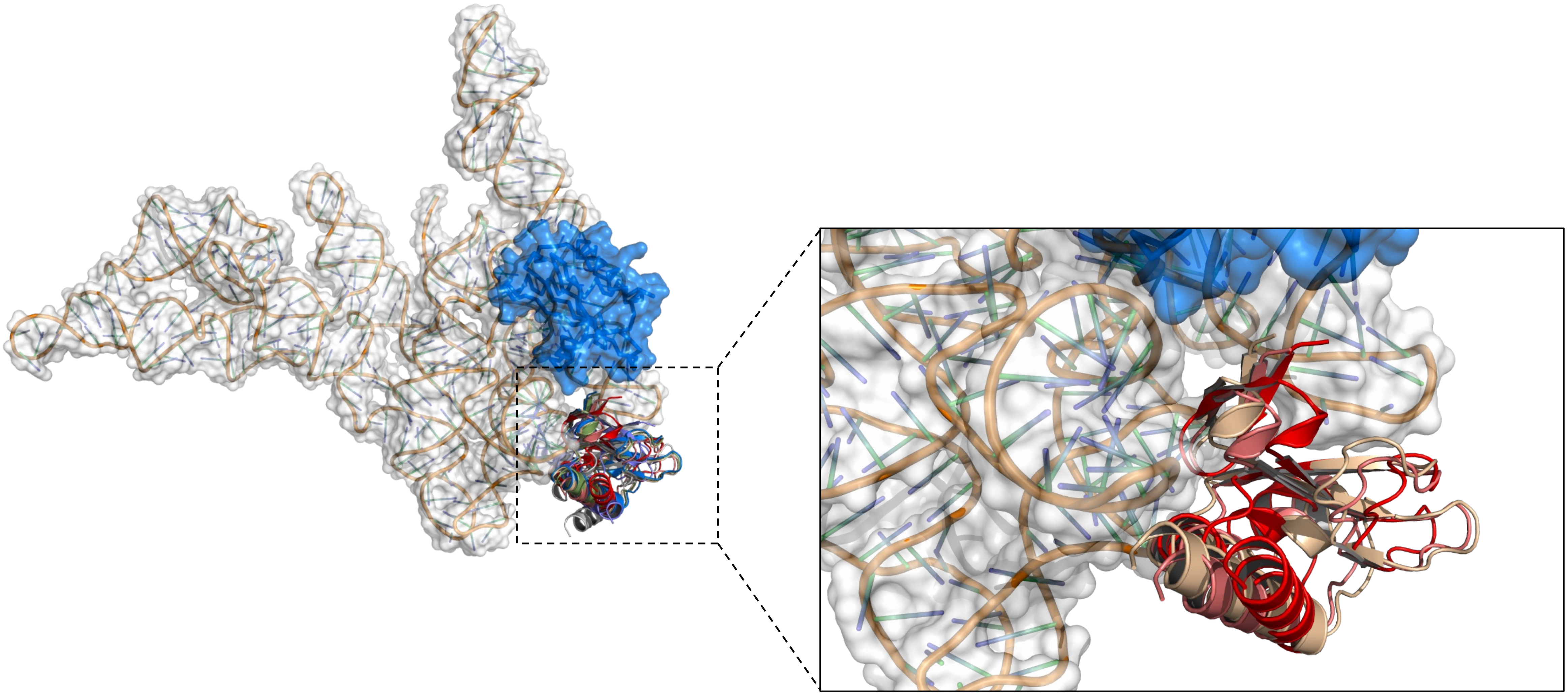

Fig S2. Accumulation of RnpM hits in the predicted P RNA binding site. Multiple RnpM decoys (cartoon representation) were predicted to bind in close proximity to the RnpA protein (transparent surface, marine).

Wicke et al., Fig. S2

**Table S1. Cross-links identified for RnpM and P RNA**

| RnpM peptide sequence | RNA-adduct | Crosslinked nucleotide | Localization score | #q-value | Crosslinker |
|---|---|---|---|---|---|
| Localization score (LS) 1.0 | | | | | |
| CVVTGEM(Oxidation)KPK | G+C4H6O2 | G | 2,641 | 0 | DEB |
| CVVTGEM(Oxidation)KPK | U+C4H6O2 | U | 1,422 | 1,15E-06 | DEB |
| CVVTGEM(Oxidation)KPK | A+C4H6O2 | A | 3,699 | 1,00E-06 | DEB |
| CVVTGEMKPK | A+C4H6O2 | A | 3,315 | 2,35E-06 | DEB |
| CVVTGEM(Oxidation)KPK | U+C4H6O2 | U | 1,181 | 1,93E-06 | DEB |
| SKEGEISVDPTGK | U+C4H6O2 | U | 1,902 | 3,30E-06 | DEB |
| SKEGEISVDPTGK | U+C4H6O2 | U | 1,826 | 2,39E-06 | DEB |
| SKEGEISVDPTGK | U+C4H6O2 | U | 1,096 | 8,78E-07 | DEB |
| KCVVTGEM(Oxidation)KPK | CG+C5H9N1 | C | 1,196 | 3,65E-06 | NM |
| CVVTGEM(Oxidation)KPK | A+C5H9N1 | A | 3,116 | 2,25E-06 | NM |
| CVVTGEM(Oxidation)KPK | U+C5H9N1 | U | 2,811 | 3,61E-06 | NM |
| CVVTGEM(Oxidation)KPK | A+C5H9N1 | A | 2,908 | 2,97E-06 | NM |
| CVVTGEM(Oxidation)KPK | G+C5H9N1 | G | 1,132 | 2,96E-06 | NM |
| CVVTGEMKPK | C+C5H9N1 | C | 1,608 | 2,10E-06 | NM |
| KCVVTGEMKPK | A+C5H9N1 | A | 1,454 | 3,87E-06 | NM |
| ECILAAK | G+C5H9N1 | G | 1,029 | 4,38E-06 | NM |
| SKEGEISVDPTGKK | G+C5H9N1 | G | 1,638 | 4,42E-06 | NM |
| SKEGEISVDPTGKK | G+C5H9N1 | G | 1,787 | 4,52E-06 | NM |
| ECILAAK | A+C5H9N1 | A | 1,647 | 4,53E-06 | NM |
| EGEISVDPTGK | G+C5H9N1 | G | 1,208 | 3,90E-06 | NM |
| CVVTGEM(Oxidation)KPK | AA+C5H9N1 | A | 1,094 | 4,30E-06 | NM |
| EGEISVDPTGKK | G+C5H9N1 | G | 2,106 | 4,09E-06 | NM |
| GAYLTLDKECILAAK | CC+C5H9N1-H3O4P1 | C | 1,166 | 8,91E-03 | NM |
| CVVTGEMKPK | U+C5H9N1 | U | 1,147 | 3,98E-06 | NM |
| ECILAAK | A+C5H9N1 | A | 1,000 | 4,82E-06 | NM |
| GAYLTLDKECILAAK | CC+C5H9N1-H3O4P1 | C | 2,010 | 9,55E-04 | NM |

| | | | | | |
|---|---|---|---|---|---|
| GAYLTLDKECILAAK | GU+C5H9N1-H1O3P1 | G | 1,229 | 1,86E-03 | NM |
| GAYLTLDKECILAAK | AU+C5H9N1-H1O3P1 | A | 1,583 | 4,97E-06 | NM |
| GAYLTLDKECILAAK | AU+C5H9N1-H1O3P1 | A | 1,040 | 2,76E-03 | NM |
| GAYLTLDKECILAAK | CC+C5H9N1-H3O4P1 | C | 1,435 | 5,36E-06 | NM |
| GAYLTLDKECILAAK | G+C5H9N1 | G | 1,187 | 4,93E-06 | NM |
| GAYLTLDKECILAAK | U+C5H9N1 | U | 1,883 | 4,48E-06 | NM |
| GAYLTLDKECILAAK | A+C5H9N1 | A | 1,848 | 2,76E-03 | NM |
| GAYLTLDKECILAAK | G+C5H9N1-H2O1 | G | 1,270 | 4,13E-06 | NM |
| GAYLTLDKECILAAK | U+C5H9N1 | U | 1,218 | 8,91E-03 | NM |
| | | LS 1.5 | | | |
| CVVTGEM(Oxidation)KPK | G+C4H6O2 | G | 2,641 | 0 | DEB |
| CVVTGEM(Oxidation)KPK | A+C4H6O2 | A | 3,699 | 1,00E-06 | DEB |
| CVVTGEMKPK | A+C4H6O2 | A | 3,315 | 2,35E-06 | DEB |
| CVVTGEM(Oxidation)KPK | U+C4H6O2 | U | 1,591 | 2,15E-06 | DEB |
| SKEGEISVDPTGK | U+C4H6O2 | U | 1,902 | 3,30E-06 | DEB |
| SKEGEISVDPTGK | U+C4H6O2 | U | 1,826 | 2,39E-06 | DEB |
| CVVTGEM(Oxidation)KPK | A+C5H9N1 | A | 3,116 | 2,25E-06 | NM |
| CVVTGEM(Oxidation)KPK | U+C5H9N1 | U | 2,811 | 3,61E-06 | NM |
| CVVTGEM(Oxidation)KPK | A+C5H9N1 | A | 2,908 | 2,97E-06 | NM |
| CVVTGEMKPK | C+C5H9N1 | C | 1,608 | 2,10E-06 | NM |
| SKEGEISVDPTGKK | G+C5H9N1 | G | 1,638 | 4,42E-06 | NM |
| SKEGEISVDPTGKK | G+C5H9N1 | G | 1,787 | 4,52E-06 | NM |
| ECILAAK | A+C5H9N1 | A | 1,647 | 4,53E-06 | NM |
| EGEISVDPTGKK | G+C5H9N1 | G | 2,106 | 4,09E-06 | NM |
| GAYLTLDKECILAAK | CC+C5H9N1-H3O4P1 | C | 2,010 | 9,55E-04 | NM |
| GAYLTLDKECILAAK | AU+C5H9N1-H1O3P1 | A | 1,583 | 4,97E-06 | NM |

| | | | | | |
|---|---|---|---|---|---|
| GAYLTLDKECILAAK | A+C5H9N1 | A | 2,929 | 3,66E-06 | NM |
| GAYLTLDKECILAAK | U+C5H9N1 | U | 1,883 | 4,48E-06 | NM |
| GAYLTLDKECILAAK | A+C5H9N1 | A | 1,848 | 2,76E-03 | NM |
| | | LS 2.0 | | | |
| CVVTGEM(Oxidation)KPK | G+C4H6O2 | G | 2,641 | 0 | DEB |
| CVVTGEM(Oxidation)KPK | A+C4H6O2 | A | 3,699 | 1,00E-06 | DEB |
| CVVTGEMKPK | A+C4H6O2 | A | 3,315 | 2,35E-06 | DEB |
| SKEGEISVDPTGK | U+C4H6O2 | U | 2,042 | 2,33E-06 | DEB |
| EGEISVDPTGK | U+C4H6O2 | U | 5,763 | 3,09E-06 | DEB |
| CVVTGEM(Oxidation)KPK | A+C5H9N1 | A | 3,116 | 2,25E-06 | NM |
| CVVTGEM(Oxidation)KPK | U+C5H9N1 | U | 2,811 | 3,61E-06 | NM |
| CVVTGEM(Oxidation)KPK | A+C5H9N1 | A | 2,908 | 2,97E-06 | NM |
| SKEGEISVDPTGKK | G+C5H9N1 | G | 3,365 | 4,93E-06 | NM |
| EGEISVDPTGKK | G+C5H9N1 | G | 2,106 | 4,09E-06 | NM |
| GAYLTLDKECILAAK | CC+C5H9N1-H3O4P1 | C | 2,010 | 9,55E-04 | NM |
| GAYLTLDKECILAAK | A+C5H9N1 | A | 2,929 | 3,66E-06 | NM |
| GAYLTLDKECILAAK | A+C5H9N1 | A | 2,474 | 3,68E-06 | NM |
| GAYLTLDKECILAAK | U+C5H9N1 | U | 2,336 | 4,73E-06 | NM |

**Table S2. Cross-links identified for RnpA and P RNA**

| RnpA peptide sequence | RNA-adduct | Crosslinked nucleotide | Localization score | #q-value | Crosslinker |
|---|---|---|---|---|---|
| Localization score (LS) 1.0 | | | | | |
| IGNAVM(Oxidation)R | U+C4H6O2 | U | 1,132 | 3,57E-06 | DEB |
| PASQLTYEETKK | U+C4H6O2 | U | 1,185 | 1,64E-06 | DEB |
| KPASQLTYEETK | G+C5H9N1 | G | 1,253 | 1,24E-06 | NM |
| KPASQLTYEETK | AU+C5H9N1 | A | 2,006 | 1,26E-06 | NM |
| KPASQLTYEETK | C+C5H9N1 | C | 1,056 | 2,43E-06 | NM |
| KPASQLTYEETK | G+C5H9N1 | G | 1,069 | 3,12E-06 | NM |
| HGTSVANR | G+C5H9N1 | G | 1,227 | 3,26E-06 | NM |
| HGTSVANR | A+C5H9N1 | A | 1,523 | 3,37E-06 | NM |
| VFKHGTSVANR | G+C5H9N1 | G | 1,112 | 3,62E-06 | NM |
| IGNAVMR | G+C5H9N1 | G | 1,598 | 3,69E-06 | NM |
| SLQHLFR | G+C5H9N1 | G | 1,614 | 4,00E-06 | NM |
| KIGNAVM(Oxidation)R | G+C5H9N1 | G | 1,219 | 4,31E-06 | NM |
| VGLSVSKK | G+C5H9N1 | G | 1,416 | 4,33E-06 | NM |
| KPASQLTYEETKK | A+C5H9N1 | A | 1,148 | 4,52E-06 | NM |
| NEDFQK | A+C5H9N1 | A | 1,775 | 4,53E-06 | NM |
| LS 1.5 | | | | | |
| IGNAVM(Oxidation)R | U+C4H6O2 | U | 1,669 | 2,54E-06 | DEB |
| KPASQLTYEETK | AU+C5H9N1 | A | 2,006 | 1,26E-06 | NM |
| HGTSVANR | A+C5H9N1 | A | 1,523 | 3,37E-06 | NM |
| IGNAVMR | G+C5H9N1 | G | 1,598 | 3,69E-06 | NM |
| SLQHLFR | G+C5H9N1 | G | 1,614 | 4,00E-06 | NM |
| NEDFQK | A+C5H9N1 | A | 1,775 | 4,53E-06 | NM |
| KPASQLTYEETK | AU+C5H9N1 | A | 2,006 | 1,26E-06 | NM |
| HGTSVANR | A+C5H9N1 | A | 2,931 | 4,03E-06 | NM |

**Table S3. RnpM decoys predicted and classified according to the localization score of cross-linking data**

| Model abbreviation | Buried surface area | Number of hydrogen bonds | d0 | d0-ALL | d2 | d2-ALL | d3 | d3-ALL | d5 | d5-All | Number of interactions | Number of short range contacts |
|---|---|---|---|---|---|---|---|---|---|---|---|---|
| Localization score (LS) 1.0 | | | | | | | | | | | | |
| model_3216 | 814,94 | 14 | 31 | 54 | 35 | 91 | 36 | 114 | 37 | 176 | 43 | 2 |
| model_4971 | 932,448 | 27 | 28 | 57 | 34 | 93 | 35 | 118 | 36 | 160 | 50 | 4 |
| model_6561 | 703,288 | 18 | 31 | 55 | 35 | 100 | 36 | 118 | 37 | 173 | 29 | 3 |
| model_1228 | 947,319 | 40 | 34 | 61 | 35 | 95 | 35 | 119 | 37 | 163 | 97 | 10 |
| model_742 | 860,192 | 24 | 31 | 57 | 35 | 96 | 37 | 119 | 37 | 164 | 45 | 1 |
| model_1268 | 886,657 | 24 | 33 | 55 | 35 | 100 | 37 | 120 | 37 | 185 | 30 | 3 |
| model_2804 | 906,434 | 24 | 32 | 56 | 35 | 99 | 37 | 121 | 37 | 186 | 31 | 4 |
| model_156 | 775,197 | 34 | 33 | 70 | 36 | 108 | 36 | 123 | 37 | 163 | 55 | 2 |
| model_3530 | 628,927 | 22 | 33 | 56 | 33 | 93 | 35 | 124 | 37 | 173 | 41 | 2 |
| model_8396 | 754,519 | 32 | 31 | 60 | 33 | 90 | 35 | 124 | 35 | 165 | 40 | 3 |
| model_2463 | 801,658 | 37 | 31 | 63 | 33 | 91 | 35 | 125 | 35 | 172 | 46 | 6 |
| model_345 | 967,083 | 32 | 32 | 55 | 35 | 100 | 36 | 125 | 37 | 174 | 53 | 4 |

| | | | | | | | | | | | | |
|---|---|---|---|---|---|---|---|---|---|---|---|---|
| model_3219 | 885,739 | 24 | 30 | 57 | 35 | 107 | 35 | 127 | 37 | 176 | 52 | 6 |
| model_214 | 905,135 | 29 | 32 | 54 | 36 | 99 | 36 | 128 | 37 | 175 | 48 | 2 |
| model_4338 | 844,206 | 28 | 30 | 55 | 34 | 108 | 36 | 131 | 37 | 182 | 47 | 7 |
| model_636 | 1064,82 | 25 | 31 | 65 | 36 | 114 | 37 | 132 | 37 | 191 | 56 | 5 |
| model_30 | 1002,58 | 35 | 33 | 59 | 36 | 107 | 36 | 137 | 37 | 190 | 67 | 2 |
| model_85 | 876,26 | 32 | 34 | 63 | 36 | 112 | 36 | 138 | 37 | 192 | 68 | 1 |
| model_4911 | 678,253 | 23 | 31 | 63 | 35 | 108 | 35 | 139 | 37 | 181 | 42 | 2 |
| model_3350 | 774,026 | 27 | 29 | 56 | 35 | 113 | 36 | 140 | 37 | 193 | 41 | 3 |
| LS 1.5 | | | | | | | | | | | | |
| model_1228 | 947,319 | 40 | 19 | 33 | 20 | 54 | 20 | 64 | 20 | 78 | 97 | 10 |
| model_742 | 860,192 | 24 | 17 | 35 | 19 | 55 | 20 | 67 | 20 | 81 | 45 | 1 |
| model_167 | 1093,14 | 49 | 17 | 34 | 19 | 60 | 20 | 69 | 20 | 99 | 62 | 6 |
| model_695 | 1145,62 | 52 | 17 | 37 | 19 | 58 | 20 | 70 | 20 | 106 | 72 | 9 |
| model_156 | 775,197 | 34 | 19 | 39 | 20 | 60 | 20 | 71 | 20 | 86 | 55 | 2 |
| model_1950 | 851,603 | 37 | 17 | 33 | 19 | 75 | 20 | 86 | 20 | 110 | 67 | 8 |
| model_926 | 780,727 | 24 | 17 | 37 | 19 | 76 | 19 | 96 | 20 | 133 | 53 | 1 |
| model_1228 | 947,319 | 40 | 19 | 33 | 20 | 54 | 20 | 64 | 20 | 78 | 97 | 10 |

| | | | | | | | | | | | | |
|---|---|---|---|---|---|---|---|---|---|---|---|---|
| LS 2.0 | | | | | | | | | | | | |
| model_742 | 860,192 | 24 | 13 | 27 | 14 | 42 | 15 | 51 | 15 | 59 | 45 | 1 |
| model_695 | 1145,62 | 52 | 13 | 30 | 14 | 44 | 15 | 53 | 15 | 79 | 72 | 9 |
| model_156 | 775,197 | 34 | 14 | 30 | 15 | 46 | 15 | 54 | 15 | 66 | 55 | 2 |

**Table S4. Best scoring decoys for the RnpM-P RNA complex**

| **Model abbreviation** | **Buried surface area** | **Number of hydrogen bonds** | **d0** | **d0-ALL** | **d2** | **d2-ALL** | **d3** | **d3-ALL** | **d5** | **d5-All** | **Number of interactions** | **Number of short range contacts** |
|---|---|---|---|---|---|---|---|---|---|---|---|---|
| model_6561 | 703,288 | 18 | 31 | 55 | 35 | 100 | 36 | 118 | 37 | 173 | 29 | 3 |
| model_1228 | 947,319 | 40 | 34 | 61 | 35 | 95 | 35 | 119 | 37 | 163 | 97 | 10 |
| model_156 | 775,197 | 34 | 33 | 70 | 36 | 108 | 36 | 123 | 37 | 163 | 55 | 2 |
| model_345 | 967,083 | 32 | 32 | 55 | 35 | 100 | 36 | 125 | 37 | 174 | 53 | 4 |
| model_214 | 905,135 | 29 | 32 | 54 | 36 | 99 | 36 | 128 | 37 | 175 | 48 | 2 |
| model_30 | 1002,58 | 35 | 33 | 59 | 36 | 107 | 36 | 137 | 37 | 190 | 67 | 2 |
| model_85 | 876,26 | 32 | 34 | 63 | 36 | 112 | 36 | 138 | 37 | 192 | 68 | 1 |
